# DIRT/μ – Automated extraction of root hair traits using combinatorial optimization

**DOI:** 10.1101/2024.01.18.576310

**Authors:** Peter Pietrzyk, Neen Phan-Udom, Chartinun Chutoe, Lise Pingault, Ankita Roy, Marc Libault, Patompong Johns Saengwilai, Alexander Bucksch

## Abstract

Similar to any microscopic appendages, such as cilia or antennae, phenotyping of root hairs has been a challenge due to their complex intersecting arrangements in two-dimensional (2D) images and the technical limitations of automated measurements. Digital Imaging of Root Traits at Microscale (DIRT/μ) addresses this issue by computationally resolving intersections and extracting individual root hairs from 2D microscopy images. This solution enables automatic and precise trait measurements of individual root hairs. DIRT/μ rigorously defines a set of rules to resolve intersecting root hairs and minimizes a newly designed cost function to combinatorically identify each root hair in the microscopy image. As a result, DIRT/μ accurately measures traits such as root hair length (RHL) distribution and root hair density (RHD), which are impractical for manual assessment. We tested DIRT/μ on three datasets to validate its performance and showcase potential applications. By measuring root hair traits in a fraction of the time manual methods require, DIRT/μ eliminates subjective biases from manual measurements. Automating individual root hair extraction accelerates phenotyping and quantifies trait variability within and among plants, creating new possibilities to characterize root hair function and their underlying genetics.

## Introduction

Phenotypes exist across scales of organismal organization and change over time. This multiscale characteristic poses a major challenge in phenomics, making a comprehensive measurement of phenotypes nearly impossible^1^. Computational tools analyzing digital images partly overcome this phenotyping problem and have been used to study the traits of human, animal, and plant phenotypes^2^. However, self-occlusions due to the 2D projection during imaging pose a major challenge for image-based measurements of phenotypes^3,4^. This issue arises not only at large spatial scales, such as the branching of a tree, but also at the finest, microscopic scales. A common, yet often small morphological trait present in many organisms is a collection of elongated appendages with important protective, physiological, or sensory functions, such as antennae, hairs, and trichomes, which are attached to the body of the organism^5^. Each appendage has a simple linear morphology that could be easily traced and measured, but collectively, these appendages intersect with one another or can completely occlude one another when imaged. As a result, and despite the ubiquitous presence of elongated appendages in nature, computational tools to extract individual elongated appendages from images are scarce, making the comprehensive phenotyping of organisms with such appendages impossible.

An example of such appendages is root hairs, which are elongated epidermal cells of the root that laterally extend into the soil^6^. Root hairs display distinguishing characteristics in length, density, and occurrence along the roots, which vary distinctively among species and genotypes and across environments. Root hair traits have been linked to higher water uptake and improved tolerance to drought^7–9^. In particular, the traits RHL and RHD have been further associated with increased tolerance to low levels of immobile nutrients in the soil. Under low levels of phosphorus (P), the traits RHL and RHD increase, and longer and denser root hairs increase P absorption^10,11^. Similar benefits have been described for low levels of mobile nutrients. Plants with longer root hairs have greater biomass under low nitrogen (N), a mobile nutrient, compared to plants with short root hairs^12,13^. Root hair development can be separated into three distinct and complex pathways: the differentiation of epidermal root cells into root hair cells, the initiation of root hair growth, and root hair elongation underlie different and complex pathways^14^. Therefore, it is unsurprising that the control of RHL and RHD (e.g., in rice) underlies separate genomic regions^15^. Due to their role in nutrient and water uptake, root hairs provide an important breeding target for sustainable agriculture under drought and increasingly nutrient-depleted soils^16,17^. To leverage the benefits of root hairs in agricultural production, the function and control of specific root hair traits must be addressed^18^.

Trait measurements of root hairs are challenging due to their small scale, three-dimensional (3D) nature and often-complex arrangement. Microscopes, in combination with digital cameras, enable the capture of images of root hairs, which can be used to measure their traits. Bright-field and dark-field light microscopes, either compound or stereo, are the most common types for imaging root hairs due to their low cost and simple operation^12,19,20^. Other microscopy techniques, including confocal, electron, and light sheet fluorescent microscopy, can increase extractable information, but due to their higher cost, they are less accessible for most studies^21–25^. Similarly, imaging technologies for 3D measurements of root hairs exist but are financially unattainable for most labs and require data acquisition times that do not allow for high through-put phenotyping^26–28^. Root hairs can be counted and traced manually in microscopy images using software, such as ImageJ^29^, to determine RHL and RHD. However, this manual task is time-consuming, tedious, and subjective to the researcher. On the other hand, determining these traits automatically is challenging due to the projection of root hairs onto a 2D plane, which causes combinatorial challenges to resolve the occlusions and intersections in the image data.

In recent years, semi-automated and automated methods have approached the combinatorial challenges with proxy measurements to phenotype root hairs. Vincent, et al. ^30^ used multivariate linear regression to determine root hair area *in situ* based on the machine-learning classification of root hair pixels in minirhizotron images. A method by Guichard, et al. ^31^ measures the length of mature root hairs, the distance of emerging root hairs to the root tip, and the length of the root hair growth zone and estimates the root hair growth rate, but this requires root hairs to grow orthogonal to the root and does not determine RHD. The results are based on slicing the profile of the root hairs along the root as a proxy measurement to quantify RHL and RHD, rather than measurements of individual root hairs. The results of this method were used to study the relevance of root hair traits for N acquisition in Arabidopsis^13,32^. A third method, by Lu, et al. ^33^, classifies root hair pixels using deep learning to measure root hair area. Measured RHL is based on the length of a skeletal line of root hair regions across the entire image but does not separate individual hairs. All three methods identify root hair pixels with high accuracy but do not obtain actual counts of root hairs that resolve the complex challenges of occluding root hairs along an imaged root segment to calculate length and density traits.

In response to the challenges regarding root hair phenotyping, we developed a method for extracting individual root hairs automatically from microscopy images in a statistically robust way. Our approach uses machine learning to classify image pixels as either root hair, root, or background because the 3D root hairs are being projected onto a 2D image plane. The projection leads to many intersections of root hairs in the imaging data whose resolution is beyond the capability of the human visual system. Resolving these many intersections in an image is non-trivial because every intersection of two or more root hairs can be resolved in multiple ways. Resolving this network of intersections can result in trillions of possible outcomes for a single image. We demonstrate that our approach works even in dense arrangements with many intersecting root hairs in a feasible time frame. As a result, we can measure the number of root hairs and the distribution of individual lengths and shapes. In this paper, we demonstrate the robustness and detail of the information derived from our method on three datasets.

## Method

The algorithm works through sequential steps that begin by identifying image pixels as root hairs and result in the detection and representation of individual root hairs (Figure 1). First, image pixels are classified to distinguish between root hair, root, and background. From the classified root hair pixels, a medial axis is extracted as a skeleton. Next, we fit weighted splines to the medial axis to obtain a smooth line representation of potential root hair segments. The core idea of DIRT/µ is then to test whether combinations of these splines adhere to a priori criteria of root hair shape to compute iteratively a set of measurable root hairs. The selected solution is the set of root hairs that minimizes the a priori criteria of root hair shape. The results enable us to compute estimations of traits, such as the number of root hairs, their shape, and their distribution.

**Figure 1:**
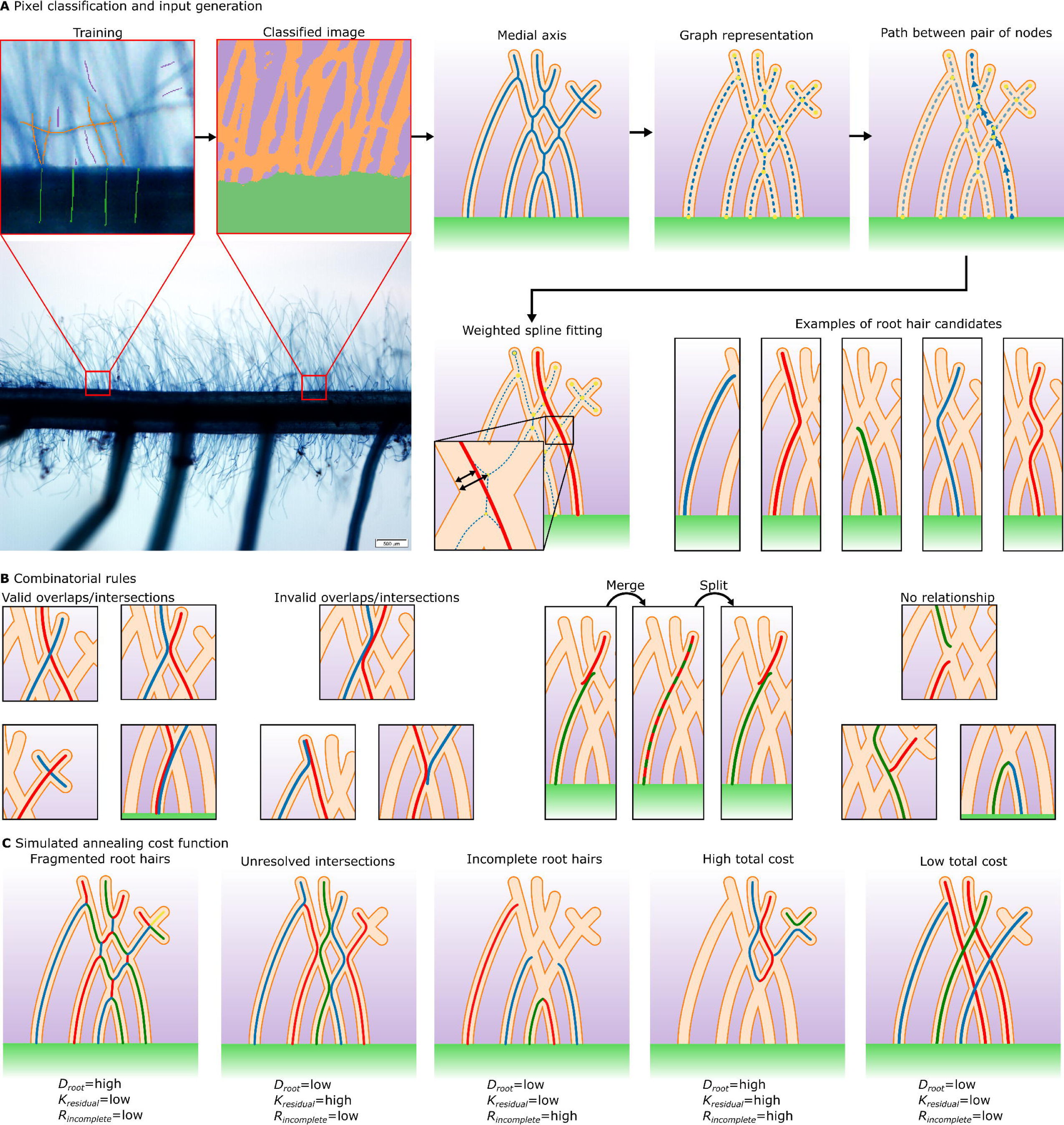
Workflow for extracting root hair from a microscopy image. (A) To extract root hairs from a microscopy image, we create a training set with classes for root hair, root, and background to classify pixels in the images. We perform a medial axis transform on classified root hair pixels, identify junctions and termination points as nodes in the medial axis, and select a path between a set of two nodes. Along the path, we fit a spline weighted by the diameter of the medial circle. We repeat the weighted spline fitting for all close-by node pairs and retain only valid splines. (B) We establish rules between pairs of splines. Splines can exist side-by-side if they intersect with large angles, touch slightly, or emerge from the same node at the root surface and overlap. Splines must not exist side-by-side if long internal sections of splines overlap, two splines overlap at their end if they emerge from the same node not at the root surface, or if an end section of a spline overlaps with the internal section of another spline. Two splines are treated as single root hairs if only two of their ends overlap. Removing one of the splines splits the root hair. (C) Combinations of root hairs are optimized by minimizing the three metrics of distance to the root, curvature, and incompleteness. Fragmented root hairs result in a high distance to the root, incorrectly resolved root hairs result in high curvatures, and incomplete root hairs result in a high cost for incompleteness.

### Ilastik classifies pixels to standardize input data across various imaging modalities

To reconstruct individual root hairs, we need to identify all pixels that correspond to root hairs. In addition, we need to identify pixels corresponding to the parent root to determine where root hairs emerge. We used the software ilastik to classify all image pixels as either root hair, root, or background^34^. The software provides a user interface to create training sets by manually annotating pixels according to their class and to preview the classification result interactively. For the classification, we use a random forest classifier with features obtained from morphological filters on multiple spatial scales (for more details, see Methods S1 in Supplementary Material). Typically, a training set can be created within a few hours, and all root hair images from the same dataset can be classified in a batch process. Small segments of root hair or root are removed from the classified image (for more details, see Methods S1 in Supplementary Material). The benefit of using this machine-learning approach to classify pixels is that datasets obtained under different lab protocols can be used as input for DIRT/µ (Figure 1A).

### Building a relational graph data structure to order root hair segments

The classified image provides only rasterized data without information to identify individual root hairs. All the classified root hairs together result in a network of inseparable regions representing root hairs. We extract the medial axis from root hair pixels to create a relational graph of individual medial axis segments (Figure 1A). Each branching of the medial axis results from touching or intersecting root hairs. Each point on the medial axis is approximated by an image pixel and belongs to one of three types: 1) a termination point, if it has only one neighboring pixel on the medial axis; 2) a medial axis segment (i.e., a line), if it has exactly two neighboring points on the medial axis; and 3) a junction point, if it has more than two neighboring medial axis points. The relational graph of medial axis segments is then computed. Nodes are embedded at the locations of the termination and junction points of the medial axis, and edges connect the nodes wherever a single medial axis segment connects termination and/or junction points. To consider parallel edges in the graph, we add one node to the relational graph, representing paths from one junction or termination point of the medial axis to the next junction or termination point. The medial axis contains information about the curvature of root hair segments, and the relational graph illustrates the neighborhood topology of adjacent segments.

### Rules to fit valid splines select root hair candidates as a combination of root hair segments

Based on the medial axis and the relational graph, it is possible to create a root hair candidate. This candidate is created by traversing a path along the relational graph from one node to another node and by fitting a smooth line to this path (Figure 1A). Traversing paths between alternative pairs of nodes and alternative paths between the same pair of nodes results in additional root hair candidates. To limit the number of candidates, only nodes within reasonable proximity are used as pairs (see Methods S1 in Supplementary Material for details). For all paths, we use a weighted spline fitting method to create smooth lines, which remove medial axis artifacts at junctions to represent root hairs more accurately (see Methods S1 in Supplementary Material for details). We accomplish this task by weighting the spline at a location using Equation 1, in which *r_MA_* is the measured radius of the medial circle, *r_min_* is the interpolated value between local minima of the measured radius along the medial axis, and 0.5 accounts for noise in the raster-based medial axis. The weights of individual spline points are increased iteratively until all points are between the root hair edge and the medial axis. As a result, spline sections at junctions with high radii are smoothed more than at thinner locations of the medial axis.

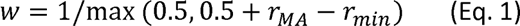

To reduce the number of splines further, we remove any invalid ones. We define an invalid spline as one with high curvature at junction points. We compute the *total curvature* for each spline and the *segment curvatures* for individual segments between adjacent nodes of the spline. *Total* and *segment curvatures* are defined as the integral of the spline curvature along its entire and segment length, respectively. At each segment, we additionally determine a reference value as the smallest possible *segment curvature* at this segment based on the *segment curvatures* of all splines overlapping with this segment. We then compare the *total curvature* of a spline with the sum of the reference values of all its segments. This process enables us to quantify the spline’s *residual spline curvature* to quantify its curviness compared with the optimal case of the lowest possible *total curvature*. Similarly, we locally compare each individual *segment curvature* of a spline with its reference value to obtain the *residual segment curvature*. Since the curviness value is scale-independent, we can define a threshold value of 180° for *residual spline curvature* and a threshold of 45° for *residual segment curvature* to remove invalid splines. As a result, only those splines remain that have low curvature at junction points of the medial axis.

### Analyzing spline relationships reduces combinatorial space

Some of the remaining splines will touch one another or overlap partially. Therefore, we define three relationships between splines and assign one relationship to each pair of splines (Figure 1B). First, splines that overlap or intersect in an invalid manner result in conflicting relationships. Splines that have a conflict are not allowed to be both parts of the same solution. Second, in all remaining cases, splines either do not touch or they intersect or overlap in a valid manner. Third, splines that overlap at exactly one end are merged into one root hair candidate. This categorization enables us to reduce the number of splines and, thus, keep the combinatorial space small.

### Combinatorial optimization identifies splines that correspond to root hairs

We use simulated annealing to find combinatorically the state regarding the set of splines that minimizes a predefined cost function to reconstruct actual root hairs correctly and completely^35^. Our method is based on solving a 0-1 integer problem with restored feasibility at each move^36^. The process begins with a feasible state of randomly selected splines (i.e., all relationships between splines are valid) and iteratively changes the state of the spline set by either removing or adding a spline. If a newly added spline violates the relationships, then conflicting splines are removed to reestablish the validity of the current state of the spline set. Similarly, splines are merged or split when necessary. This method ensures that all intersecting root hairs are resolved validly. Furthermore, at each iteration, the cost of the current state of the spline set is re-evaluated. By decreasing the probability of accepting worse states during the simulated annealing process, the cost is forced to decrease, such that a global optimum is found at the end of the annealing process. The parametrization of the simulated annealing process is automated and is based on the description by Orsila, et al. ^37^ (see Methods S1 in Supplementary Material for detailed description). DIRT/µ quantifies the cost of a state of the spline set (i.e., how good it is) as a function of three parameters (Figure 1C, Table 1).

1. *Minimum distance to root D_root_*: We minimize the distances between each root hair candidate and the root in a least squares sense to minimize outliers at large distances to the root.
2. *Residual total curvature Κ_residual_*: We minimize the residual spline curvature of all candidates in a least squares sense to avoid candidates with excessive curvature.
3. *Incompleteness R_incomplete_*: We minimize the ratio between the length of all unresolved medial axis segments (*l_unresolved MA_)* and the total length of the medial *axis (l_MA_)* to quantify the degree of identified root hairs.

**Table 1:** Overview of optimization metrics. The table contains the calculation for the minimum distance to root, residual total curvature, and incompleteness, as well as their normalized values and explanations. The total cost is the weighted root-mean- square value of their normalized values.

|  | Values | Normalized values | Explanations |
| --- | --- | --- | --- |
| *Minimum distance to root* | $D_{\text{root}} = \sqrt{\frac{1}{n} \sum_{i=1}^n d_i^2}$ | $\tilde{D}_{\text{root}} = \frac{D_{\text{root}} - 1}{D_{\text{max}} - 1}$ | $n$ is number of candidates; $d_i$ is distance of candidate $i$ to root in pixels; $D_{\text{max}}$ equals avg. length of all medial axis segments that connect to root. |
| *Residual total curvature* | $K_{\text{residual}} = \sqrt{\frac{1}{n} \sum_{i=1}^n \kappa_i^2}$ | $\tilde{K}_{\text{residual}} = \frac{K_{\text{residual}}}{K_{\text{max}}}$ | $n$ is number of candidates; $\kappa_i$ is residual spline curvature of candidate $i$; $K_{\text{max}}$ equals avg. of all residual total curvatures. |
| *Incompleteness* | $R_{\text{incomplete}} = 1 - \frac{l_{\text{unresolved MA}}}{l_{\text{MA}}}$ | $\tilde{R}_{\text{incomplete}} = R_{\text{incomplete}}$ | $l_{\text{MA}}$ is length of all medial axis segments; $l_{\text{unresolved MA}}$ is the length of medial axis segments that are unresolved. |
| *Total cost* | $\text{cost} = \sqrt{w_1 \tilde{D}_{\text{root}}^2 + w_2 \tilde{K}_{\text{residual}}^2 + w_3 \tilde{R}_{\text{incomplete}}^2}, \sum_{k=1}^3 w_k = 1$ | | $w_1$, $w_2$, and $w_3$ are user specified weights (see Figs. S7 and S8 for sensitivity analysis of weights). |

The *total cost* of the state of the spline set is computed as the weighted root-mean-square of all three metrics to balance their minimization. In conjunction with the three types of relationships between splines, the combinatorial optimization enables us to minimize the *total* cost and, thus, resolve intersecting root hairs in a computationally efficient manner. As a result, not all splines will perfectly fit the criteria, but the number of incorrectly identified root hairs is minimized, enabling the statistical evaluation of individual root hair measurements. Therefore, we use the state with the lowest *total cost* as our solution.

### Root hair traits are measured relative to identified root hair splines

Given the identified root hairs in the state with the lowest *total cost*, we measure the lengths of all root hairs and the number of root hairs. We distinguish between root hairs on the main root (i.e., the root with largest diameter) and on thinner secondary roots emerging from the main root. Based on the length of the top and bottom edges of the main root and the number of root hairs, we determine RHD across the entire root, the densities on the top and bottom edges, the highest density in a 1 mm section, and the variation of density along the root. Further parameters can be derived from splines that represent root hairs; for example, parameters based on the curvature of the spline, which quantifies the rate of change in direction of the longitudinal axis of the root hair, such as the integral of absolute curvature along its length, average curvature, and maximum curvature.

### Three test cases demonstrate the accuracy, precision, and reliability of DIRT/μ

We used root hair images from three experiments to demonstrate the feasibility of DIRT/µ and its benefits over manual root hair measurements (Table 2). Dataset Mahidol I consists of 15 selected images of rice (*Oryza sativa*), maize (*Zea mays*), and common bean (*Phaseolus vulgaris*) roots and serves as a validation set. In this set, the images were chosen to represent the phenotypic variation, ranging from low to high RHDs, as well as short to long root hairs per species. We performed image classification in ilastik on each image individually due to differences in magnification between the images and to decouple the effects of the classification from root hair extraction. In all 15 images, we manually traced all visible root hairs and registered their lengths for comparison with the automatic measurements in ImageJ. Dataset Mahidol II, which is similar in experimental setup to Dataset Mahidol I, serves as a demonstration set. A total of 731 images were taken from rice, maize, and common bean roots, with three genotypes for each species, three to four samples per genotype, and in most samples between four and six images taken along the root. A total of 46 images, in which more than 50% of the root length was mislabeled as lateral root (e.g., due to being images not taken according to DIRT/µ protocol or due to the incorrect classification of pixels), were excluded after visual inspection. In contrast to the first dataset, we manually measured the length of only five representative root hairs and density along a representative section (approx. 1mm) of the root. Dataset University of Nebraska-Lincoln (UNL), differing from the other datasets regarding experimental setup and image acquisition, is a subset of a large dataset in which extended depth of field (EDF) images and validation data are available. This dataset served as an additional validation set, using sorghum (*Sorghum bicolor*) in well-watered and drought-stressed conditions. After removing outliers through visual inspection (30 images) and excluding additional images in which fewer than two root hairs were detected or manually measured (97 images), 359 measurement points were available.

**Table 1:** Summary of datasets used to evaluate the trait measurements of DIRT/μ.

|  | Dataset Mahidol I | Dataset Mahidol II | Dataset UNL |
| --- | --- | --- | --- |
| Number of images (plus images not analyzed) | 15 (0) | 685 (46) | 359 (127) |
| Species (# genotypes, # plants per genotype, # images per plant) | Rice (1,5,1),<br>Maize (1,5,1),<br>Common bean (1,5,1) | Rice (3, 4, 5–6),<br>Maize (3, 3–4, 1–7),<br>Common bean (3, 3–4, 4–7) | Sorghum: 32 genotypes in drought, 31 genotypes in well-watered |
| Treatments | Control, -N, -P, -K | Control, -N, -P, -K | Well-watered, drought |
| Manual measurements | All root hairs | Length: 5 representative root hairs<br>Density: representative section (appr. 1mm) | All root hairs and up to 10 or 20 in some images |
| Microscope | Olympus MVX10 | Olympus MVX10 | Leica MZ10 F |
| Color space | RGB | RGB | Gray / monochrome |
| Staining | Toluidine blue | Toluidine blue | Trypan blue |
| EDF | No | No | Yes |
| Resolution in $\mu\text{m}/\text{pixel}$ | 0.34–1.34 | Rice: 0.67, maize: 1.34–1.72, common bean: 1.08 | NA |
| Training set/classification | By individual image | Batch by genotype | Batch |
| Number of iterations | 1000 | 1000 | 1000 |

Using the Mahidol I data, we tested the optimization parameters *number of temperature levels* (see Methods S1 in Supplementary Material for definition) and *weights* of individual cost criteria (*w_1_*, *w_2_,* and *w_3_*) for their sensitivity and to determine their optimal values. We processed all 15 images and determined the resulting mean RHL (RHL_mean_), maximum RHL (RHL_max_), and standard deviation of RHL (RHL_std_), as well as root hair number (RH_num_) for a range of both optimization parameters. Regarding the *number of temperature levels,* we additionally determined the *total cost* of the simulated annealing process and computation time. The *number of temperature levels* ranged between 25 and 10,000 and computations were replicated five times at each value. To test *weights*, we varied individual weights between 0.02 and 0.96 under the constraint that they sum to 1.0. For each combination of *weights*, we computed the Pearson correlation coefficient and root-mean-square error (RMSE) of RHL_mean_, RHL_max_, RHL_std_, and RH_num_ between manual and automatic measurements.

Furthermore, we validated the DIRT/µ results using manual measurements of RHL statistics and RH_num_. RHL statistics and RH_num_ were computed for each individual image in Datasets Mahidol I and UNL. For RHL statistics, we computed RHL_mean_, RHL_max_, and RHL_std_. For each individual dataset, we calculated the correlation between the measurements performed manually and those taken with DIRT/µ.

For the Mahidol II data, we compared the traits RHL_mean_, RHL_std_, the coefficient of variation of RHL (RHL_CV_), and RHD between genotypes for the control group. For RHD, we took the quotient of the number of root hairs and the total length of the root edge. We further computed the correlation between measurements performed manually and with DIRT/µ by image in the same manner as for Datasets Mahidol I and UNL. For the computation of correlations, 28 images with fewer than two root hairs in either manual or automatic measurements were omitted. For the rice genotype Sungyod, we analyzed the effect of treatments RHL_mean_, RHL_std_, RHL_CV_, and RHD. For the same genotype, we further computed RHL_mean_ and RHD at different levels of organization: 1) separately for the top and bottom root edges per image, 2) per image, and 3) per plant sample (i.e., all images along the root axis). Values were obtained by pooling all the identified individual root hairs and root edge lengths at the corresponding level. Thus, we calculated RHL_mean_ for each sample as the mean of root hairs extracted from all images of the corresponding sample. We further calculated the RHD of a sample as the quotient of the number of root hairs in all sample images by the total length of root edges in all sample images. Values at the treatment and genotype level (e.g., RHL_mean_), were calculated from the results at the plant sample level. To keep all results comparable, we only accounted for root hairs on the primary root and ignored root hairs emerging from thinner secondary lateral roots.

## Results

### Sensitivity analysis revealed a rapid convergence of DIRT/ μ to optimal results

The sensitivity analysis revealed that, at low *numbers of temperature levels*, all the measured variables (RHL_mean_, RHL_max_, RHL_std_, RH_num_, and *total cost*) diverged only slightly from values at 10,000 *temperature levels* (see Figures S1–S5). Although the results suggest a smaller number is sufficient to converge toward a low *total cost* and to distinguish traits between our samples, we chose to use 1,000 *temperature levels* in our further analysis as a tradeoff to reduce computational time (see Figure S6) and to increase precision. Furthermore, varying the *weights* of individual cost criteria revealed the correlation between manual and automatic measurements of RHL_mean_ and RH_num_ was consistently high (r^2^=0.88−0.96 and r^2^=0.61−0.71, respectively) for all combinations of weights, but it was still highest for balanced weights (see Figure S7). Measurements of RMSE ranged between 44 and 94.7 µm for RHL_mean_ and between 40.6 and 52 root hairs for RH_num_. Both measurements, however, were lowest for low weights of the curvature metric and highest for low weights of the incompleteness metric (see Figure S8). As a result, we used equal values for all weights for all further analysis.

### Correlation analysis validates DIRT/μ and quantifies shortcomings in manual measurement methods

The distributions of manually and automatically measured RHLs display a good resemblance to each other (A). With 1,000 *temperature levels* and balanced weights, RHL_mean_ correlated very strongly between manual and automatic measurements in Dataset Mahidol I (r^2^=0.95, p<.001) and strongly in Dataset UNL (r^2^=0.73, p<.001) but are slightly biased and underestimated. For RHL_max_, we obtained even stronger correlations in Dataset Mahidol I (r^2^=0.97, p<.001) but the same correlation in Dataset UNL (r^2^=0.73, p<.001). The correlation in RHL_std_ was strong in Dataset Mahidol I (r^2^=0.95, p<.001) but only moderate in Dataset UNL (r^2^=0.59, p<.001). The performance regarding RH_num_ was weaker than for RHL statistics, with strong correlations in Dataset Mahidol I (r^2^=0.69, p<.001) and moderate correlations in Dataset UNL (r^2^=0.44, p<.001). The algorithm appears to undercount the number of root hairs at higher densities (see outlier in the right graph of Fig. 2B, corresponding to image BK1.3BR), and thus underestimates the number of root hairs at such densities.

**Figure 2:**
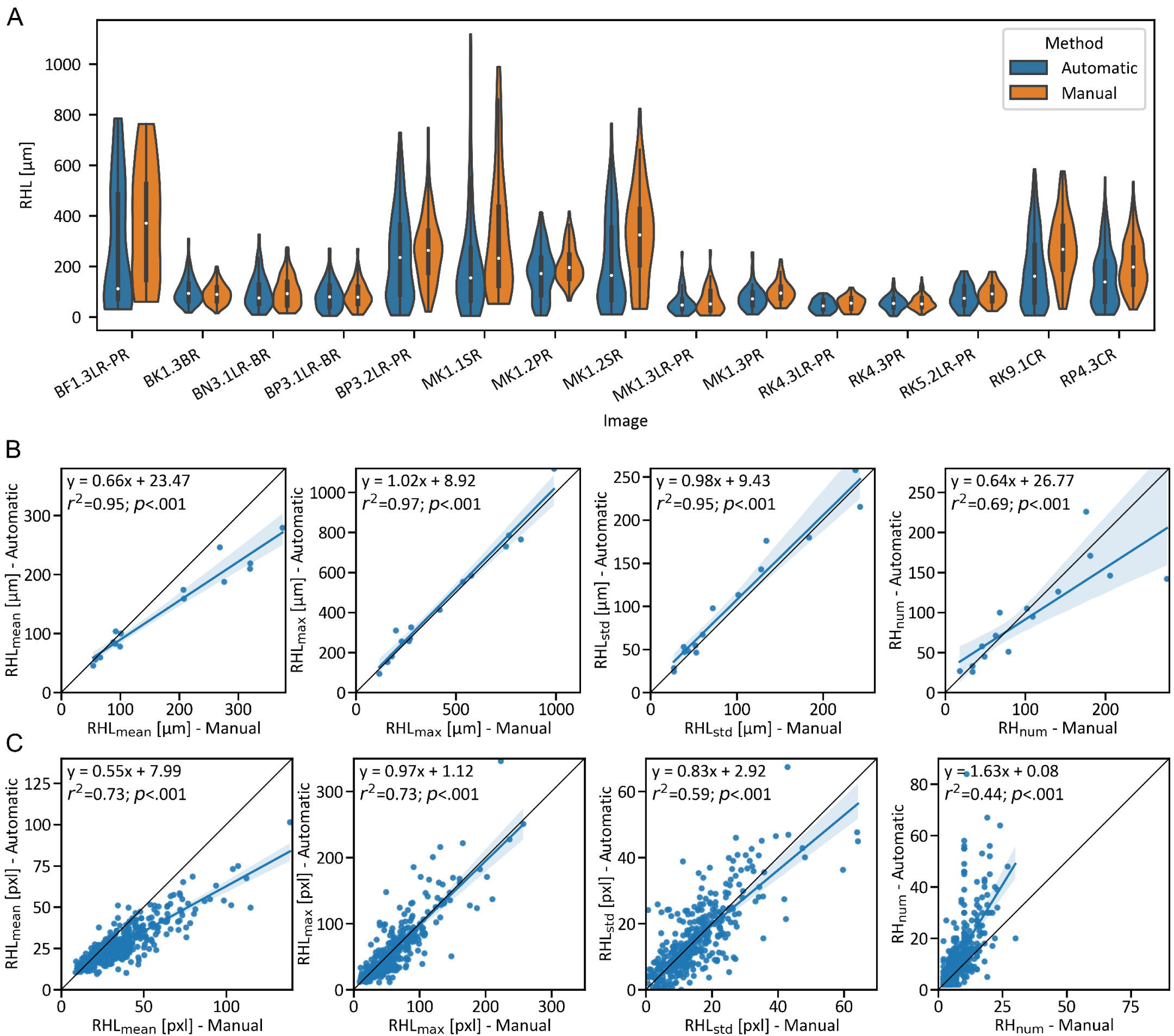
Validation of root hair measurements from automatic detection. (A) Distributions of length of all root hairs in individual images in Dataset Mahidol I from our method (blue) and manual measurements (orange). The white circle represents the median of the distribution, the black bar displays the quartiles, and the thin black centerline extends to 1.5 times the interquartile range (IQR) from the lower and upper quartiles. Each pair of automatic and manual distributions corresponds to one image. The names on the horizontal axis refer to their corresponding filenames in the supplemental material for Dataset Mahidol I. (B) Correlations of RHL_mean_, RHL_max_, RHL_std_, and RH_num_ in Dataset Mahidol I between our method (vertical axis) and manual measurements (horizontal axis). (C) Correlations of RHL_mean_, RHL_max_, RHL_std_, and RH_num_ in Dataset UNL between our method (vertical axis) and manual measurements (horizontal axis).

The measurements of RHL_mean_ in the control group of Dateset Mahidol II indicate the shortest root hairs were in rice (KDML105: 90.7 µm; Phitsanulok 2: 84.8 µm; Sungyod: 71.3 µm), followed by common bean (DOR364: 196.0 µm; L8857: 223.8 µm; SEQ7: 234.4 µm), and maize had the longest root hairs (Takfa 1: 316.8 µm; Takfa 2: 366.4 µm; Takfa 3: 450.3 µm; Figure 3A), which agreed with our expectations and manual measurements. We observed the same pattern in RHL_std_ within individual plants (Figure S11A). Using the coefficient of variation RHL_CV_, we decoupled the variation in RHL from its absolute value, revealing the smallest values in RHL_CV_ for DOR364 (0.62) and SEQ7 (0.66) and the largest value for Takfa 3 (0.83; Figure S11B). Visual inspection of these genotypes confirmed that beans generally had more uniform RHLs, and Takfa 3 had the most variation. We further observed the highest RHD in the rice genotypes KDML105 (34.04 per mm) and Phitsanulok 2 (31.02 per mm) and the smallest RHD in Takfa 1 (15.16 per mm; Figure S11C).

**Figure 3:**
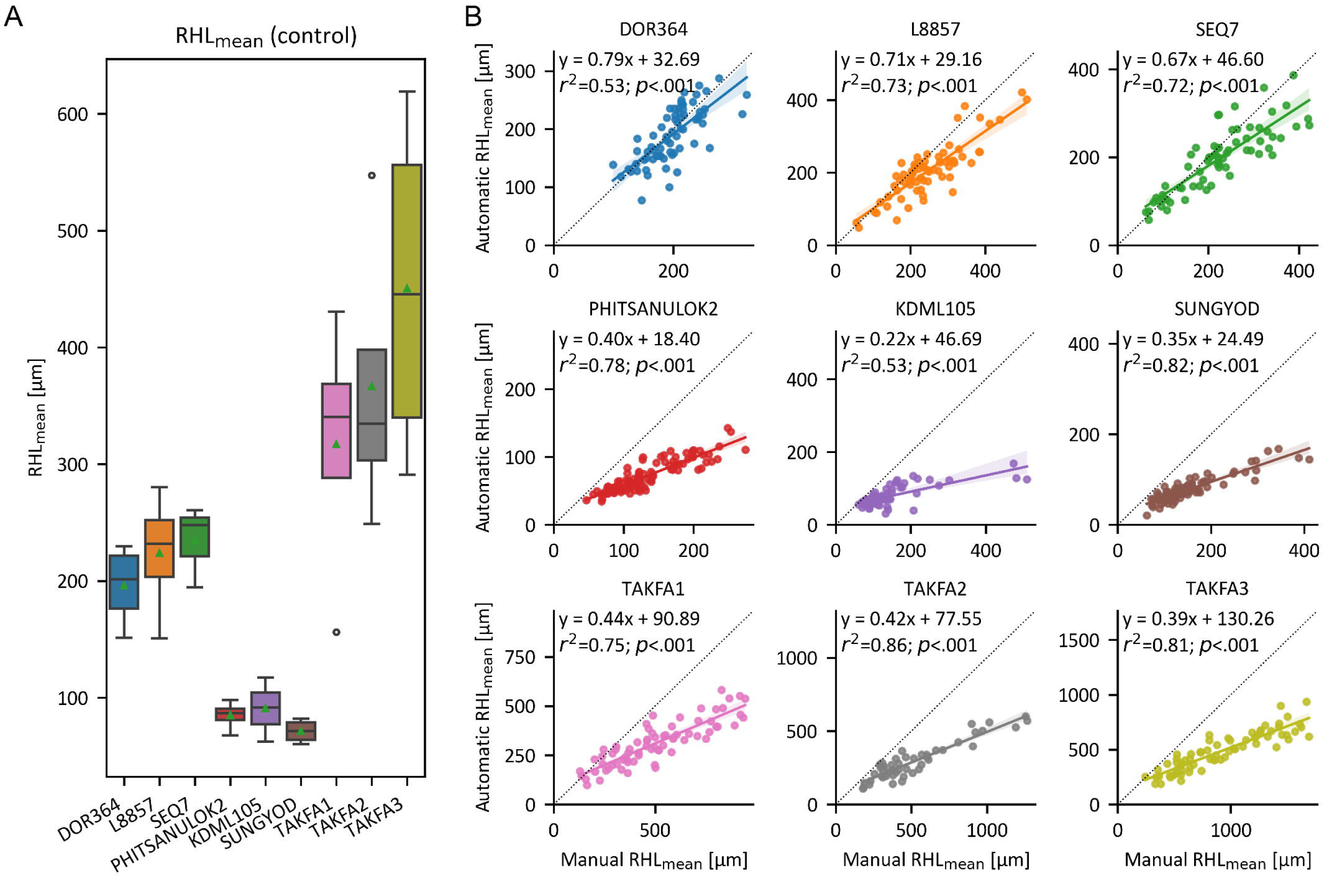
Measurements of RHL_mean_ in Dataset Mahidol II. (A) Distribution of RHL_mean_ in the control group illustrated by genotype. RHL_mean_ was computed for individual plants. The box displays the quartiles of the distribution, the middle horizontal line represents the median, and the whiskers extend to 1.5 times the IQR from the lower and upper quartiles. Outliers outside the whiskers are represented as circles. The mean of the distribution is displayed as a green triangle. (B) Correlation for automatic and manual measurements of RHL_mean_. Each plot includes the results of a single genotype. Each circle represents the RHL_mean_ per image. The lines represent the linear regression models with 95% confidence intervals.

**Figure 4:**
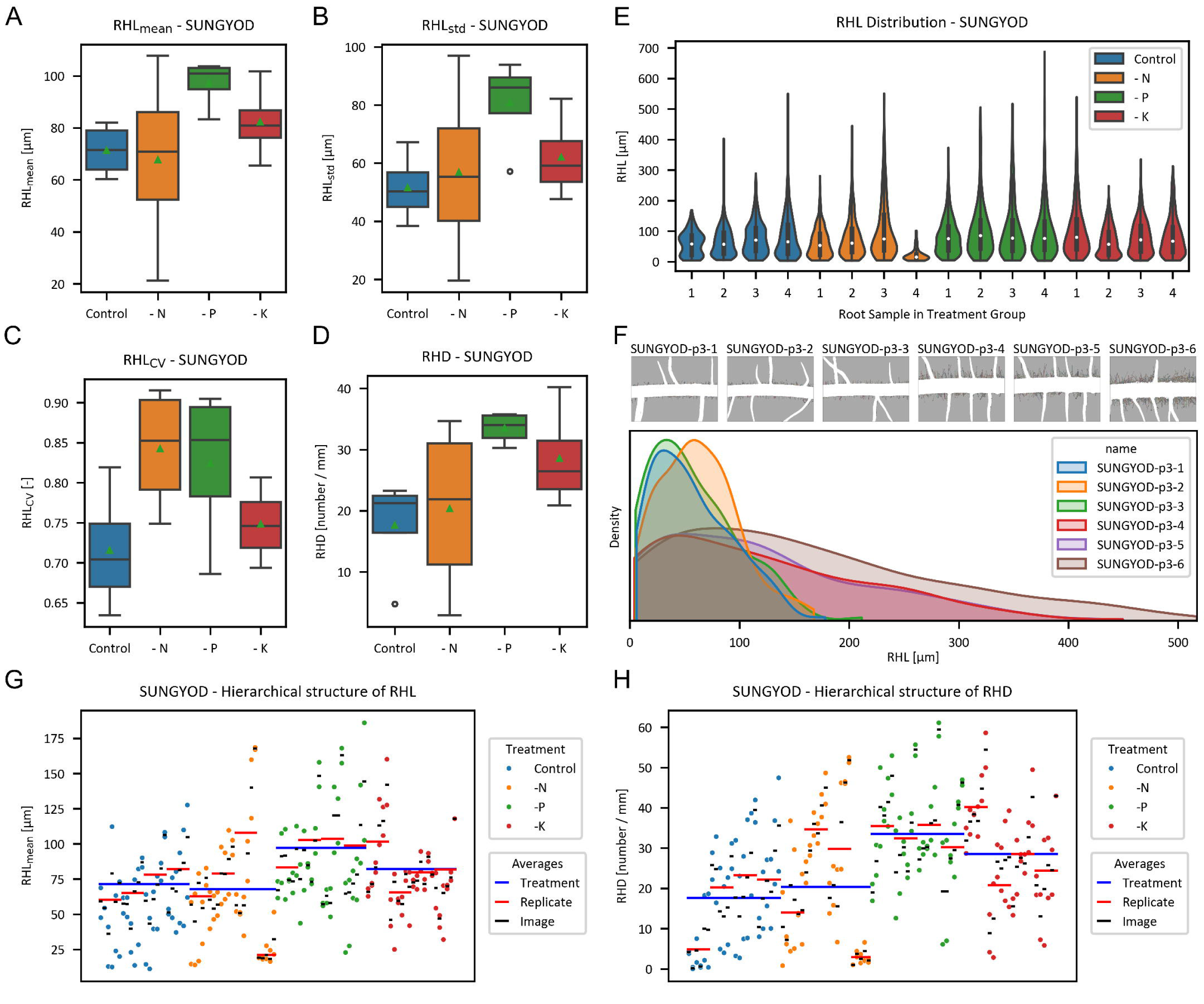
Analysis of rice genotype Sungyod in Dataset Mahidol II. (A–D) RHL_mean_ (A), RHL_std_ (B), RHL_CV_ (C) and RHD (D) were measured for all individual plants in the control, reduced nitrogen, reduced phosphorus, and reduced potassium groups. The box displays the quartiles of the distribution, the middle horizontal line represents the median, and the whiskers extend to 1.5 times the IQR from the lower and upper quartiles. Outliers outside the whiskers are represented as circles. The mean of the distributio is displayed as a green triangle. E) Distributions of length of all root hairs in individual plants colored by treatment group. The white circle represents the median of the distribution, the black bar displays the quartiles, and the thin black centerline extends to 1.5 times the IQR from the lower and upper quartiles. F) Distributions of length of all root hairs in all six individual images of Replicate 3 under phosphorus stress. G–H) Overview of RHL_mean_ (G) and RHD (H) measured at different levels of observation. The colored dots represent the measurements taken on one side of the root in a single image. A pair of colored dots aligned vertically represents one image. The corresponding black horizontal bar illustrates the measurement taken at the whole image level. Red horizontal bars represent the measurement taken from all images corresponding to a single plant. Blue horizontal lines display the mean value of all plants in each treatment group.

In Dataset Mahidol II, we compared the traits measured with DIRT/µ with those measured manually in individual images and grouped by genotype. Automatic measurements of RHL_mean_ correlated well with the manual measurements within all nine genotypes (from *r^2^*=0.53 to r*^2^*=0.86; Figure 3). The slopes of fitted regression lines were closest to one in the bean genotypes (0.67-0.79), and lowest for rice (0.22 to 0.40) and maize (0.39−0.44) genotypes. Correlations in RHL_std_ were the lowest, with values between *r^2^*=0.13 for DOR364 and *r^2^*=0.54 for Sungyod (Figure S12). Nearly all the RHL_std_ values were higher in the automatic measurements than in the manual measurements. Correlations for the RHL_max_ range from *r^2^*=0.21 for DOR364 to *r^2^*=0.76 for Phitsanulok 2 (Figure S13). Manual and automatic measurements of RHD correlated slightly weaker than measurements of RHL, ranging between *r^2^*=0.21 and *r^2^*=0.67, with bean genotypes and Takfa 2 having the lowest correlations (Figure S14). Our measurements of RHD were lower than the manual measurements in bean (slope 0.21−0.51) and maize (slope 0.34−0.61) but higher in rice (slope 1.15–1.42).

We further illustrate root hair trait measurements in the example of Sungyod (). Root hairs under phosphorus stress were 25.82 µm longer than root hairs in the control group, but despite the large difference, it was not significant (p=0.103). Results for RHL_std_ were higher in the reduced phosphorus group than in the control (p=0.028). The RHL_CV_ under reduced nitrogen and reduced phosphorus was higher than in the control group, but the difference was not significant (p=0.063 and p=0.138). The RHD under reduced phosphorus was 15.86 root hairs/mm lower than in the control group (p=0.013). The measurements using DIRT/µ resulted in the full distribution of RHL, revealing an overall pattern and small nuances, such as skewness, kurtosis, or the proportions of smaller and longer root hairs, at the plant level (, E) or the individual image level (, F). There can be differences not only between treatments but also along the root axis of a single root (, F). In general, we frequently observed differences with a factor larger than two for RHL_mean_ and RHD at all levels of observation. For example, large differences occurred between the lower and upper edges of the root in a single image, between images along the root axis, and between individual samples (, G and H).

### The degree of occlusions drives the computation time of DIRT/μ

The computation time of our method is driven by the number of junction points on the initially extracted medial axis. For Dataset Mahidol II, the computation time varied between 67 sec for KDML105-p1-1 with 24 nodes and 112.9 hours for TAKFA3-p2-6, which had 8692 nodes. The computation time for all 685 images of Dataset Mahidol II was 3408.6 hours. Combinatorial optimization accounted for most of the overall processing time (82.5%), followed by determining candidate root hairs (16.8%). Computational time increased linearly for images with more than 200 junctions in the medial axis (Figure S9), was shortest for rice images, and was longest for maize images (Figure S10).

## Discussion

### DIRT/μ eliminates user bias from root hair measurements and enables biological discovery by introducing previously inaccessible traits

Resolving intersections of root hairs in digital microscopy images enabled us to measure root hair traits that could not previously be automatically measured^38^. By extracting individual root hairs (Figure 5), we determined the distribution of RHL in each image, with parameters such as its mean, maximum, standard deviation, and coefficient of variation, as well as RHD. These traits cannot be measured manually in a feasible time because of the often long and dense root hair arrangements. Assuming a measurement time of 10 sec per root hair, it would require more than 330 hours to track manually all 119,785 root hairs that were measured automatically using DIRT/μ in Dataset Mahidol II. As such, a sub- sample of root hairs is often extracted manually to determine mean RHL and RHD. Manual measurements based on a small fraction of root hairs can result in biased (i.e., often longer and denser) root hair traits. We demonstrated that results computed with our method correspond better to manual measurements when all root hairs, rather than a subset, are measured, suggesting that we eliminate user bias.

**Figure 5:**
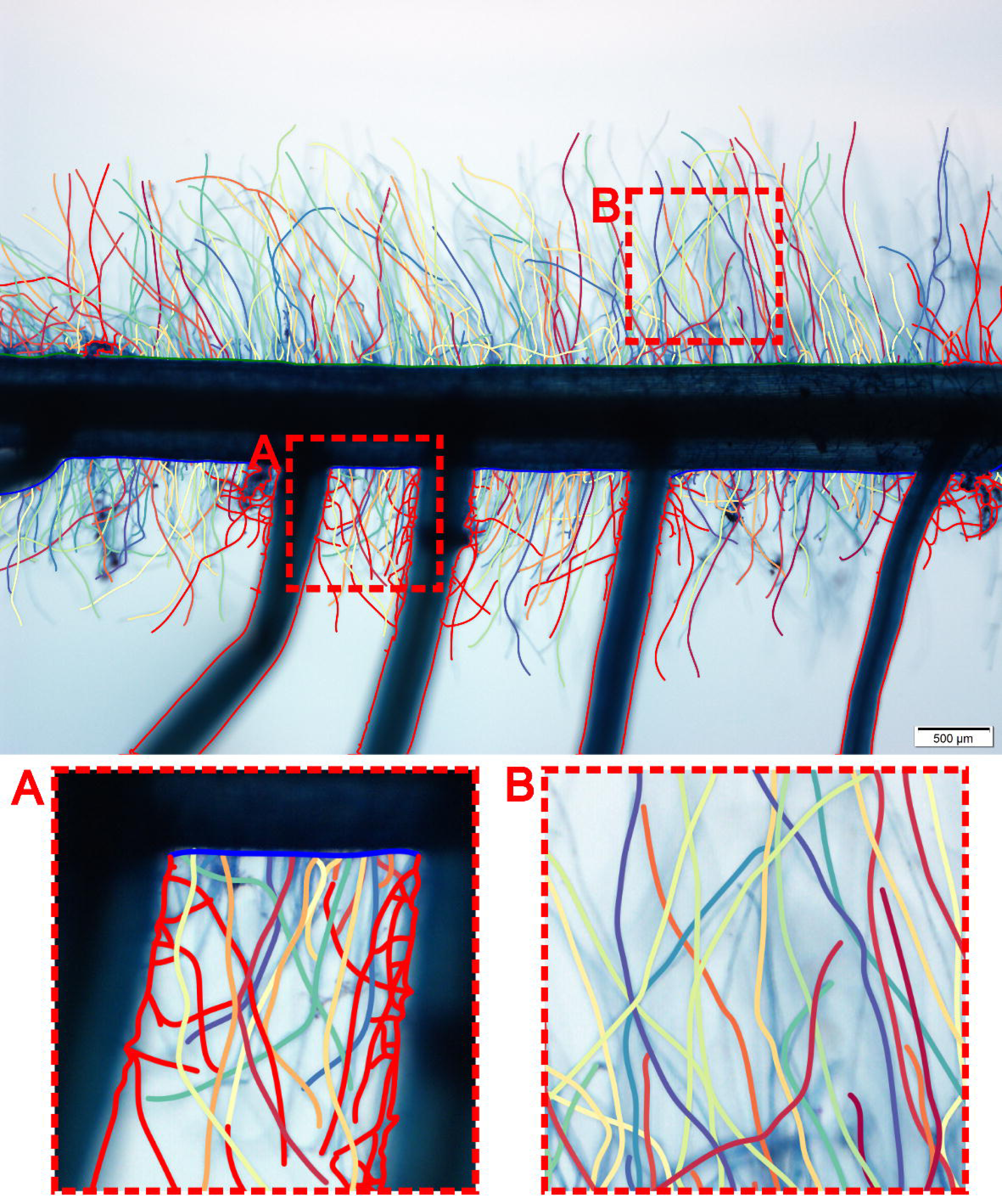
Original microscopy image of root TAKFA3-n3-1 with extracted root hairs. Valid root hairs growing on main root are plotted as lines in random colors, while root hair on lateral roots is plotted in red. The top and bottom edges of the root are plotted in green and blue, respectively. The edges of lateral roots are plotted in red. Boxes A and B show zoom-in views of extracted root hairs.

By extracting individual root hairs, our method also overcomes the limitations of previous computational methods, which were limited to determining either root hair pixel area, root hair profile, or total RHL along a section of the root^30,31,33^. From our experience, using a small focal depth minimized the remaining effects of the 2D projection on RHL and RHD measurements, such that the automatic extraction of individual root hairs from microscopy images has the potential for new biological discoveries in the plant sciences.

### DIRT/μ is an open-source solution to automated high-throughput phenotyping of root hairs

Automating the process of individual root hair extraction increases the throughput of phenotyping root traits, resulting in quicker analysis. Our measurements found a high amount of variation in root hair traits within individual plants and between plants within the same experimental groups. DIRT/μ can address this variation by increasing the number of images taken along the root and by increasing the number of samples. The higher throughput at different sampling levels and precision can improve statistical inferences in root hair research^39–41^. We note, however, that our pipeline is not fully automated, because obtaining the image classification as an input with ilastik requires human input. Nonetheless, data-specific training sets enable a diverse pool of experimental and technical setups, such as different microscopes and magnifications. Ilastik provides a simple interface, so anybody can learn to create a training set and classify a large set of microscopy images in batch processing on personal computers in a feasible time. However, unsupervised classification is an evolving field of research that promises to resolve manual training-set generation^42^. Further processing of classified images cannot be performed on personal computers and relies on high-performance computing (HPC) due to the great combinatorial complexity of resolving intersecting root hairs. Outsourcing this step to HPC through PlantIT^43^ therefore enables large-scale screening of root hairs for users without technical background.

DIRT/μ provides high-quality and affordable root hair phenotyping with good availability, accessibility, and applicability for all research labs around the world. DIRT/μ is available on GitHub^44^ and as a Docker container^45^. PlantIT and CyVerse are easily accessible to researchers without any programming knowledge^46^. Online workshops are available for end users for training through PhenomeForce^47^.

### DIRT/μ is an algorithmic basis that extends to many biological applications

Although we developed DIRT/μ primarily for, and tested it on, microscopy images, we believe it works on images taken *in situ* with mini- and micro-rhizotrons. Measurements of plant morphology across scales have great potential in plant sciences^48,49^. DIRT/μ has the potential to phenotype any hair-like structure, such as trichomes in plants, antennae or hairs in insects, or cilia in the lungs. The main principles of the DIRT/μ algorithm (i.e., combinatorial rules, cost function, and combinatorial optimization) are dimension-free, such that measurements could be extended to 3D by fitting splines to an underlying 3D skeleton. Our algorithm was developed to extract microscopic, hair-like structures; however, modification of the combinatorial rules and cost function to resolve intersecting branches while preserving branching orders means it can be adapted to identify individual branches in higher-level branching structures, such as roots, leaves, and trees. Therefore, DIRT/μ can result in more rigorous phenotyping.

## Acknowledgments

We thank Wes Bonelli for creating a Docker container and integrating DIRT/µ into PlantIT and Benjamin Smith for sharing his macro to preprocess the UNL data set. The research was supported by the NSF CAREER Award No. 1845760 and an iPlant/CyVerse Seed Grant “High-throughput Computing Platform for Quantifying Root Traits from Images” to A.B. and by NSF Award No. 2127485 to M.L. Any opinions, findings, and conclusions or recommendations expressed in this material are those of the author(s) and do not necessarily reflect the views of the National Science Foundation. The work was conducted while transitioning institutions for A.B. (University of Georgia to University of Arizona) and M.L. (University of Nebraska to University of Missouri).

## Contributions

P.P. and A.B. conceived algorithms, validation experiments and data analysis and wrote the original manuscript. P.P. implemented algorithms and performed data analysis. N.P-U., C.C., and P.J.S generated Datasets Mahidol I and II. and measured manual ground truth data, L.P. and M.L. generated Dataset UNL and measured manual ground truth data. A.R. provided parts of the literature review in the background section and performed a beta test of the software. All authors reviewed and edited the manuscript. A.B. conceived the project and acquired funding.

## Notes

https://figshare.com/s/67b1c8acab2ff8ea6fd1

### Competing Interest Statement

The authors have declared no competing interest.

