## Supplemental Material File 1 for "DIRT/μ – Automated extraction of root hair traits using combinatorial optimization"

Supplementary Material

### 1. Methods S1

**Plant material and growth conditions (Mahidol I and Mahidol II).** The plants used in the Mahidol I dataset were Kaw Dawk Mali 105 rice (KDML105), maize (commercial super sweet corn #00252), and common bean (DOR364). For the Mahidol II dataset, the rice genotypes were KDML105, Phitsanulok 2, and Sungyod, which are all Thai rice varieties. KDML105, commonly known as jasmine rice, is lowland, photoperiod-sensitive, and the most important commercial rice variety in Thailand. Phitsanulok 2 is an upland rice variety that is photoperiod-insensitive. Sungyod is a variety local to Phattalung Province in the south of Thailand that produces distinct red-colored grains. The maize genotypes used were Takfa 1, Takfa 2, and Takfa 3, all of which are inbred lines from the Nakhon Sawan Field Crops Research Center, Thailand. For common bean, the genotypes used were the high-yielding variety DOR364, the recombinant inbred line L8857 (a cross of phosphorous deficiency and drought-tolerant parents), and the drought-tolerant variety SEQ7. The rice, maize, and common bean seeds were surface-sterilized using 0.5% NaOCl for 5 mins, 6% NaOCl for 15 mins, and 50% NaOCl for 1 min, respectively. The seeds were germinated using a roll-up system, and the rolls were soaked in 0.5 mM CaSO_4_ to promote root growth. The seeds were left to germinate in complete darkness until the first shoot appeared over the roll (approximately four to seven days). The seedlings were moved to light three days before transplanting them to hydroponics culture.

The plants were grown in mason jars wrapped with tinfoil and filled with nutrient solution in 2/3 of the jar (approximately 130 mL). The full-strength (X1) control nutrient solution contained 6 mM KNO_3_, 4 mM Ca(NO_3_)_2_·10H_2_O, 2 mM NH_4_H_2_PO_4_, 1 mM MgSO_4_·7 H_2_O, and 1 mM of micronutrient stock solution. The micronutrient stock solution contained 50 µmol KCl, 25 µmol H_3_Bo_3_, 2 µmol MnSO_4_·H_2_O, 2 µmol ZnSO4·7 H_2_O, 0.5 µmol CuSO4·5 H_2_O, 0.5 µmol (NH_4_)_6_Mo_7_O_24_, and 50 µmol Fe-EDTA. The nutrient solution for each nutrient deficiency treatment was made by omitting the salts containing the nutrient in the control solution. Supplementary salts were added to maintain the concentration of other elements. The pH was adjusted to 6.5, and the nutrient solution was sterilized by autoclaving. To acclimate the plants, a quarter-strength (X1/4) nutrient solution was used in the first week and increased to half-strength (X1/2) in the second week and until the end of the experiment. The nutrient solutions were changed every three days. The maize and rice were grown in the greenhouse at Mahidol University, Salaya Campus (13°47’34.2”N 100°19’23.5”E) for 14 days after transplanting (DAP) in July and November 2019, with an average temperature of 32±3°C and 28±2°C, respectively. Common bean plants were grown for 30 DAP on a hydroponic shelf at 27±2°C with a photoperiod of 12 hours.

The shoot was cut and oven-dried at 60°C for 72 hours to obtain the shoot dry weight. The root was washed using distilled water and preserved in 70% ethanol before root hair imaging.

**Collection of ground truth (Mahidol I and II).** For the Mahidol I dataset, root segments were collected from different root classes, including primary roots from all plant species, basal roots from beans, seminal roots from maize, and crown roots from rice. Only the primary root was selected for the Mahidol II dataset. The root segment was cut from the whole root and washed with distilled water. The root was stained by dipping it in 0.5% (w/v) toluidine blue for approximately 10 sec and floating it in a Petri dish filled with water. The root hairs were observed and photographed at 0.25x to 6.3x magnification using an Olympus MVX10 microscope mounted with a camera (Olympus, Tokyo, Japan). In most samples between four and six images of root hairs were taken in the zone of maturation (starting approximately 1 cm below the base of the root). Small, soft brushes were used to untangle and straighten root hairs that clumped together and to brush off any debris that blocked the view of the root hairs. Root hair lengths and RHD were manually measured using the software ImageJ (1.52a). In Dataset Mahidol I the RHL of all root hairs was measured and all root hairs were counted. Regarding RHL in Dataset Mahidol II, only five representative root hairs were measured per image and RHD was measured as the number of root hairs per 1 mm section.

**Plant material and growth conditions (UNL).** Sorghum recombinant inbred lines (RILs) derived from parental lines RTx7000 and BTx642 were used in this experiment. Eighteen seeds of each sorghum RIL were surface-sterilized and grown in a pot containing an autoclaved mix of Vermiculite:Perlite (ratio 3:1) for five days, as described by Pingault, et al. ^49^. The seedlings for each RIL with good germination were then divided into two groups with the same number of plants. Each group was transferred into an ultrasound aeroponic system (UAS) and grown under well-water (WW) control or treatment (drought; DR) conditions at a 16 hr light : 8 hr dark cycle^49,50^. After transfer into the UASs, the plants were grown for four weeks with the same settings as the nutritive solution and delivered using a timer with the settings: 30sec ON/30sec OFF^49,51^. Then, the plants were grown for an additional two weeks under control conditions (no change in the growing settings, timer: 30sec ON/30sec OFF) for one UAS, or under drought conditions (decrease of the solution delivery, 30sec ON, and decrease the OFF time by 10sec/day until 130sec) for the other UAS. After growing for six weeks in the UASs, the root system of each plant was collected, and the lateral roots were separated from the primary root, kept in 70% ethanol on Petri dishes, and stored at -20 °C for microscopic analysis. For this analysis, the lateral roots were stained with 0.01% trypan blue for 5 min. Then, the stained roots were placed in a clean Petri dish under a microscope to locate the root hairs’ location and remove the “top” and “bottom” portions of the root that contained no visible root hairs with a scalpel. The remaining sections of roots were cut into three segments (Section 1 was the topmost portion of the root, and Section 3 was the lowermost section of the root) of equal length and placed into Petri dishes containing pure water for 15 min. Once the 15-min period had elapsed, each section was placed into the 0.01% Trypan blue staining solution for 10 min, rinsed with deionized water for 3 min, and the root section was dried with a paper towel until all visible liquid was removed. The root section was fixed on a microscope slide and placed under a Leica MZ10 F stereomicroscope with a QIMaging camera to record a 20-sec film by moving the coarse focus knob ¼ clockwise. Each film was converted into an image file using a macro in ImageJ, and the image file was used to measure the RHL using ImageJ.

**Image preparation.** We used the *Enhance Contrast* function in ImageJ (Version 1.52p) to increase homogeneity in brightness between all images in Datasets Mahidol II and UNL. For all images, we enhanced the contrast by normalizing the histogram with 2.5% of all pixels being saturated.

**Training and classification of images.** We used ilastik (Version 1.3.2) to create the training data manually and used its random forest classifier for pixel classification. All six features available in ilastik were used (i.e., Gaussian smoothing, Laplacian of Gaussian, Gaussian Gradient Magnitude, Difference of Gaussians, Structure Tensor Eigenvalues, and Hessian of Gaussian Eigenvalues). For Mahidol I, we used 0.7, 1.0, 1.6, 3.5, 5.0, 10.0 as the values for sigma at each red, green, and blue (RGB) channel to account for features at multiple scales. For Gaussian smoothing, we used an additional sigma of 0.3. For Mahidol II, we used 0.7, 1.0, 1.6, 3.5, 5.0, 10.0, 15.0, 20.0, and 50.0 for sigma and an additional sigma of 0.3 for Gaussian smoothing at each RGB channel. For the UNL dataset, we used the same settings as for Mahidol I, with the exception that we used only the available monochrome channel.

The 15 images in Dataset Mahidol I were trained and classified individually. Classes were exported into .tiff files. For Mahidol II, each genotype was trained and classified individually. Here, we used a set of 10 images for each genotype. After classification, we exported the classification probability of each class into a .tiff file and applied a Gaussian smoothing filter with a sigma equal to 1.5 to each classification probability using a Python script. Subsequently, the class with the highest probability was selected for each pixel and exported into a .tiff file with the resulting classes. This additional step resulted in more spatially homogenous classifications and a reduction of noise in the downstream process, such as the closing of tiny gaps, and improved extraction of the medial axis. For UNL, we proceeded similarly to Mahidol I and created a single training set from 47 images and extracted class probabilities for all images in a single batch process.

**Preprocessing classified images.** The DIRT/µ algorithm retains only the largest connected component of the root, removes small root hair clusters (number of pixels < square of median root hair diameter) and root hair clusters that are >10 pixels from the root, closes small gaps with morphological closing, and removes small, isolated clusters with morphological opening in root hair, root, and background.

**Medial axis extraction.** Extracting the medial axis from root hair pixels frequently results in artifacts at the transitions from root hairs to the root. Therefore, the medial axis is extracted from the union of root hair and root pixels and then clipped at the transition from root hair to root. Small branches of the medial axis are pruned if they only marginally contribute to the shape of the root hair.

**Creating candidate root hairs.** It is unfeasible to create an exhaustive set of candidate root hairs by considering all topologically possible paths on the graph. We limit the set to paths between pairs of nodes that satisfy certain criteria. The start node *s* and the end node *e* must be in the same connected graph, and the shortest distance from *s* to *e* must be 10 nodes at most. Then, all paths between *s* and *e* are determined whose length is smaller than or equal to the shortest distance plus one node.

**Spline fitting and evaluation.** The splines are fitted using the *splprep* function in SciPy^52^, such that the condition in Equation 2 is satisfied, in which *w* are the weights, *y* contains the locations of the medial axis pixels, and *g* is the corresponding location of the fitted spline. The smoothing factor *s* is set equal to the total number of points on the spline (equaling the number of points of the medial axis).

$\sum{(w\cdot(y-g))}^{2}\leq s$	(Eq. 2)

**Total curvature.** The local curvature of splines is approximated numerically using Equation 3.

$\kappa=\frac{\left| x^{'}y^{''}-y^{'}x^{''} \right|}{{(x^{'}x^{'}+y^{'}y^{'})}^{3/2}}$	(Eq. 3)

The total curvature of the spline or segment is calculated by integrating Equation 3 over the *lengths* of the spline or segment, respectively (Equation 4).

$\kappa_{\mathrm{total}}=\int\kappa ds$	(Eq. 4)

**Conflicting root hairs.** Two splines intersect in an invalid manner if, at the intersecting segments, more than 50 points of a spline are closer than √2 pixels from the other spline.

**Simulated annealing.** We use a *geometric temperature schedule* and a *normalized exponential form* for the *acceptance function*. The targeted *number of temperature levels* is user-defined, and the *number of iterations at each level* equals the number of candidate splines. As a normalization factor for the *acceptance function,* we use the *initial cost*, which is calculated as the mean cost during a simulation with a predefined number of random iterations (0.2 × *number of temperature levels* × *number of iterations at each level*). The *initial temperature* is determined through heating. Here, we use an acceptance probability of 0.95 and compute the average increase in cost from all moves of the simulation, in which the *total cost* increases. For our *final temperature,* we use an acceptance probability of 0.01 at each move, such that at each *temperature level* the acceptance probability is approximated by 0.01/*number of iterations at each level*. Based on the *initial* and *final temperatures* and the user-defined *number of temperature levels,* we can determine the *geometric temperature scaling factor* for the *geometric temperature schedule*.

Moves are defined as selecting a single random spline and either adding it to or removing it from the current state. If the spline does not exist in the current state, we try to add it based on the acceptance function, and vice versa, if the spline exists already in the current state, we try to remove it based on the acceptance function. If the move is rejected, we reestablish the previous state and attempt a new random move. The optimization process is terminated when the temperature is below the *final temperature* and sufficient consecutive moves, equaling the *number of iterations at each level*, have been rejected.

### 2. Sensitivity Analysis

#### 2.1 Temperature levels

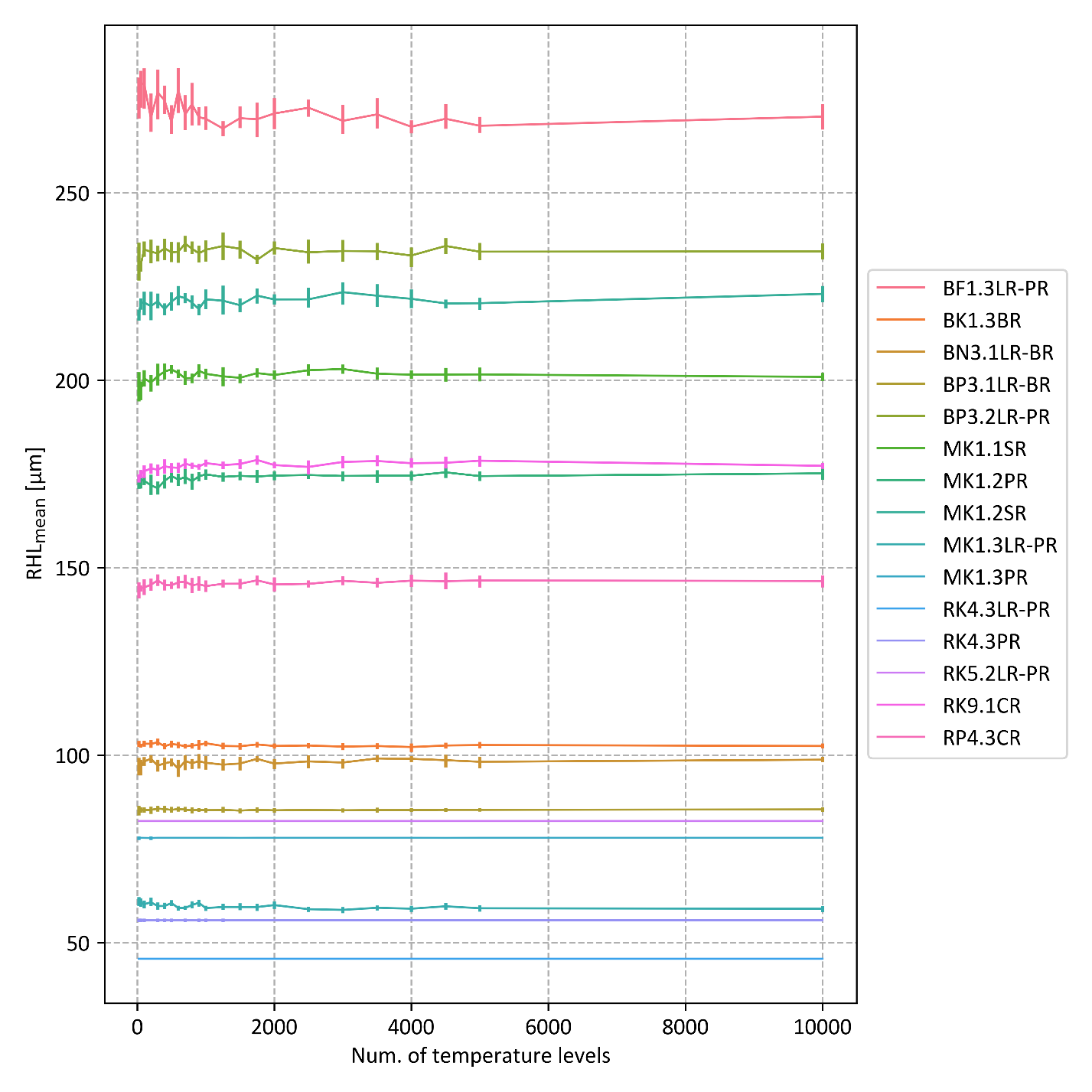

Figure S1: RHL_mean_ as a function of temperature levels for all fifteen images in Dataset Mahidol I. Each colored line represents results for one image. Measurements were taken for 25, 50, 100, 200, 300, 400, 500, 600, 700, 800, 900, 1000, 1250, 1500, 1750, 2000, 2500, 3000, 3500, 4000, 4500, 5000, and 10000 temperature levels. Results were computed five times for each image and at each level. The line indicates the corresponding mean values and the vertical bars the 95% confidence interval.

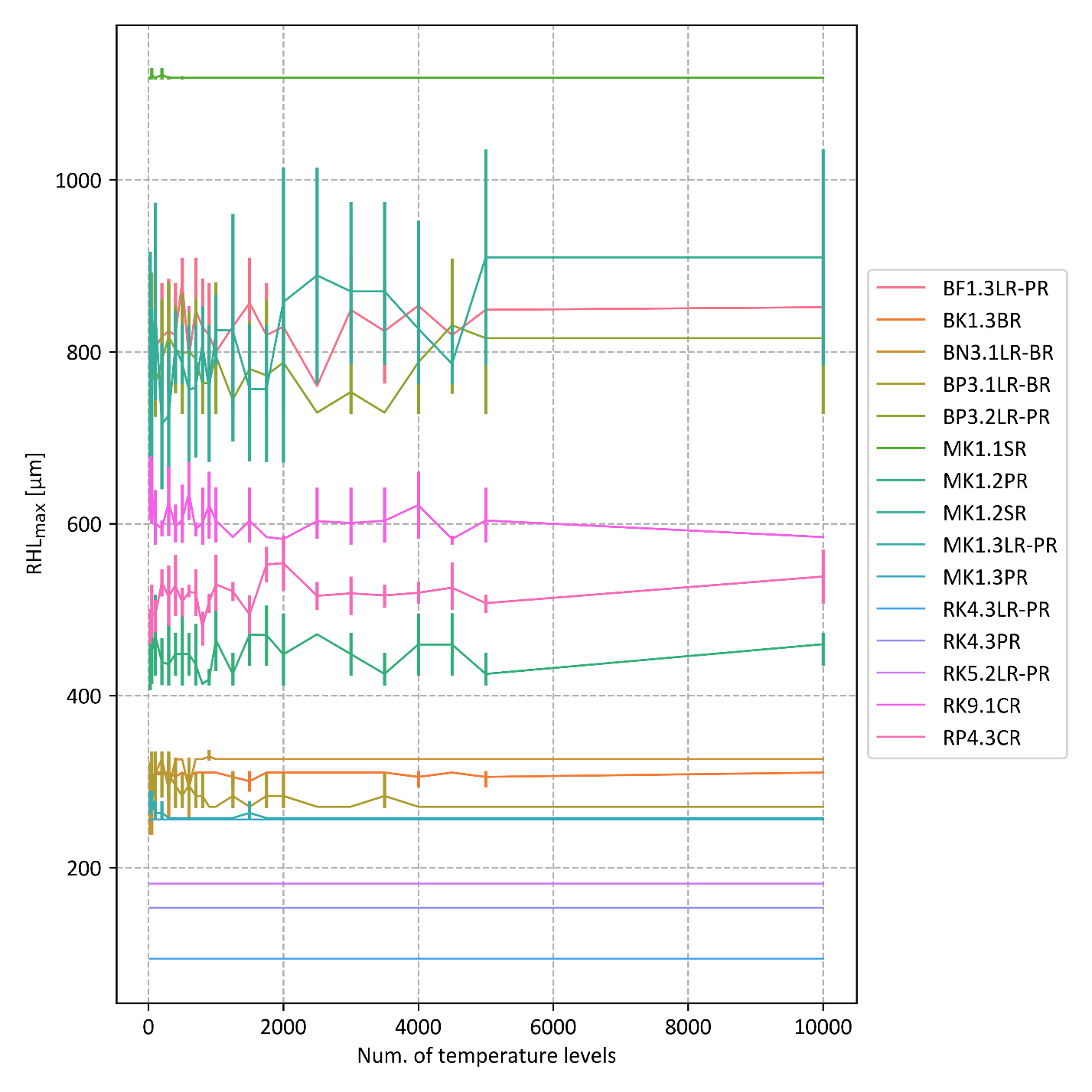

Figure S2: RHL_max_ as a function of temperature levels for all fifteen images in Dataset Mahidol I. Each colored line represents results for one image. Measurements were taken for 25, 50, 100, 200, 300, 400, 500, 600, 700, 800, 900, 1000, 1250, 1500, 1750, 2000, 2500, 3000, 3500, 4000, 4500, 5000, and 10000 temperature levels. Results were computed five times for each image and at each level. The line indicates the corresponding mean values and the vertical bars the 95% confidence interval.

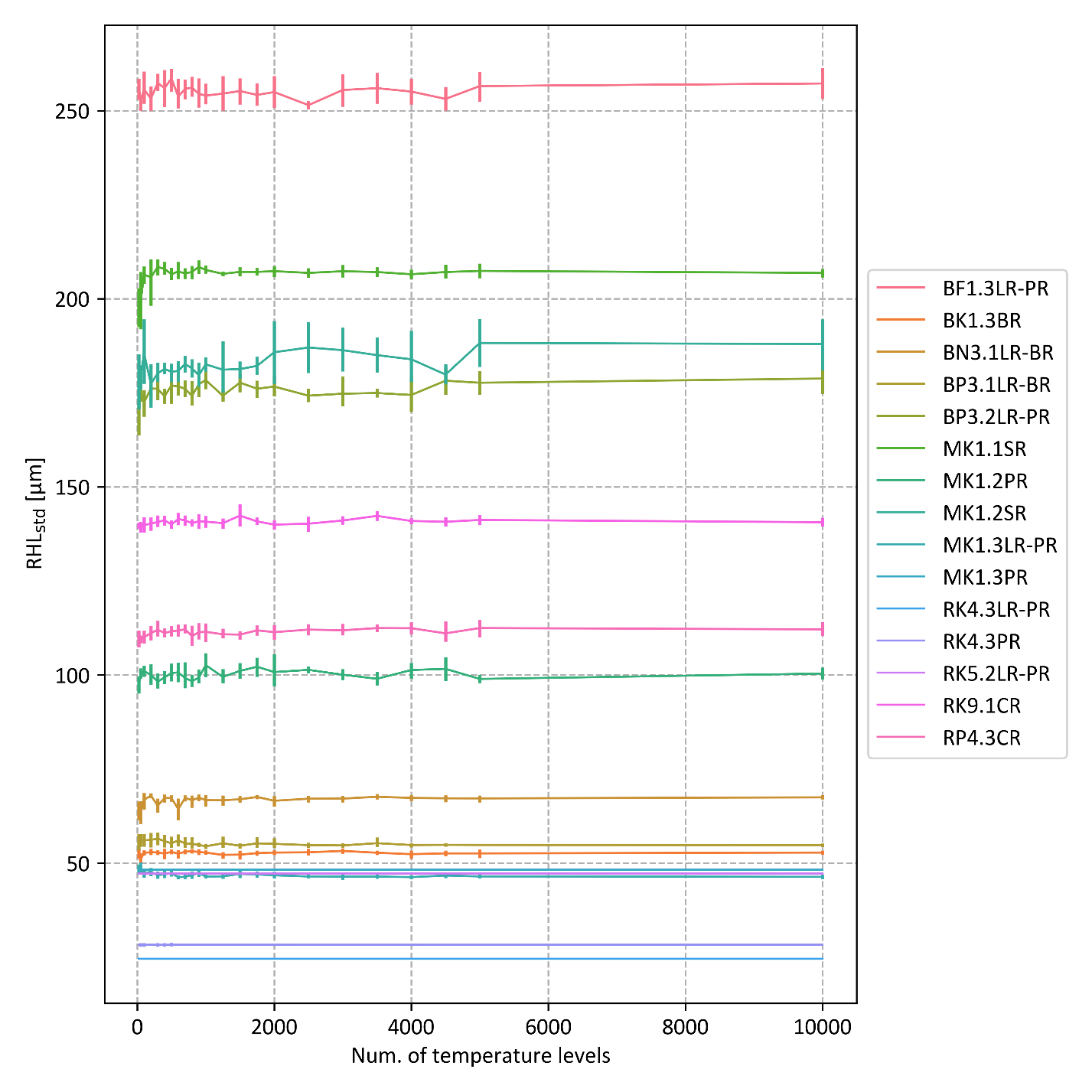

Figure S3: RHL_std_ as a function of temperature levels for all fifteen images in Dataset Mahidol I. Each colored line represents results for one image. Measurements were taken for 25, 50, 100, 200, 300, 400, 500, 600, 700, 800, 900, 1000, 1250, 1500, 1750, 2000, 2500, 3000, 3500, 4000, 4500, 5000, and 10000 temperature levels. Results were computed five times for each image and at each level. The line indicates the corresponding mean values and the vertical bars the 95% confidence interval.

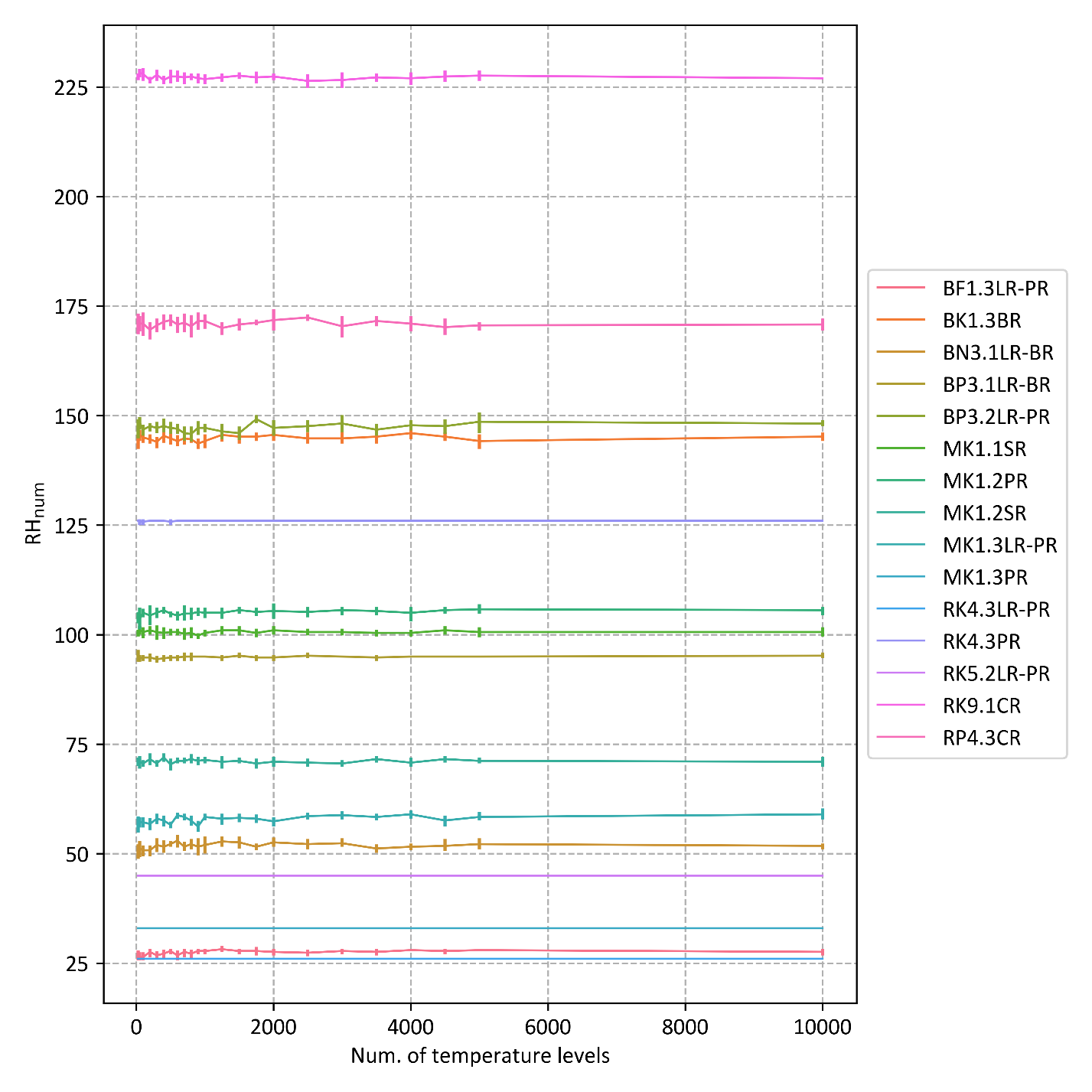

Figure S4: RH_num_ as a function of temperature levels for all fifteen images in Dataset Mahidol I. Each colored line represents results for one image. Measurements were taken for 25, 50, 100, 200, 300, 400, 500, 600, 700, 800, 900, 1000, 1250, 1500, 1750, 2000, 2500, 3000, 3500, 4000, 4500, 5000, and 10000 temperature levels. Results were computed five times for each image and at each level. The line indicates the corresponding mean values and the vertical bars the 95% confidence interval.

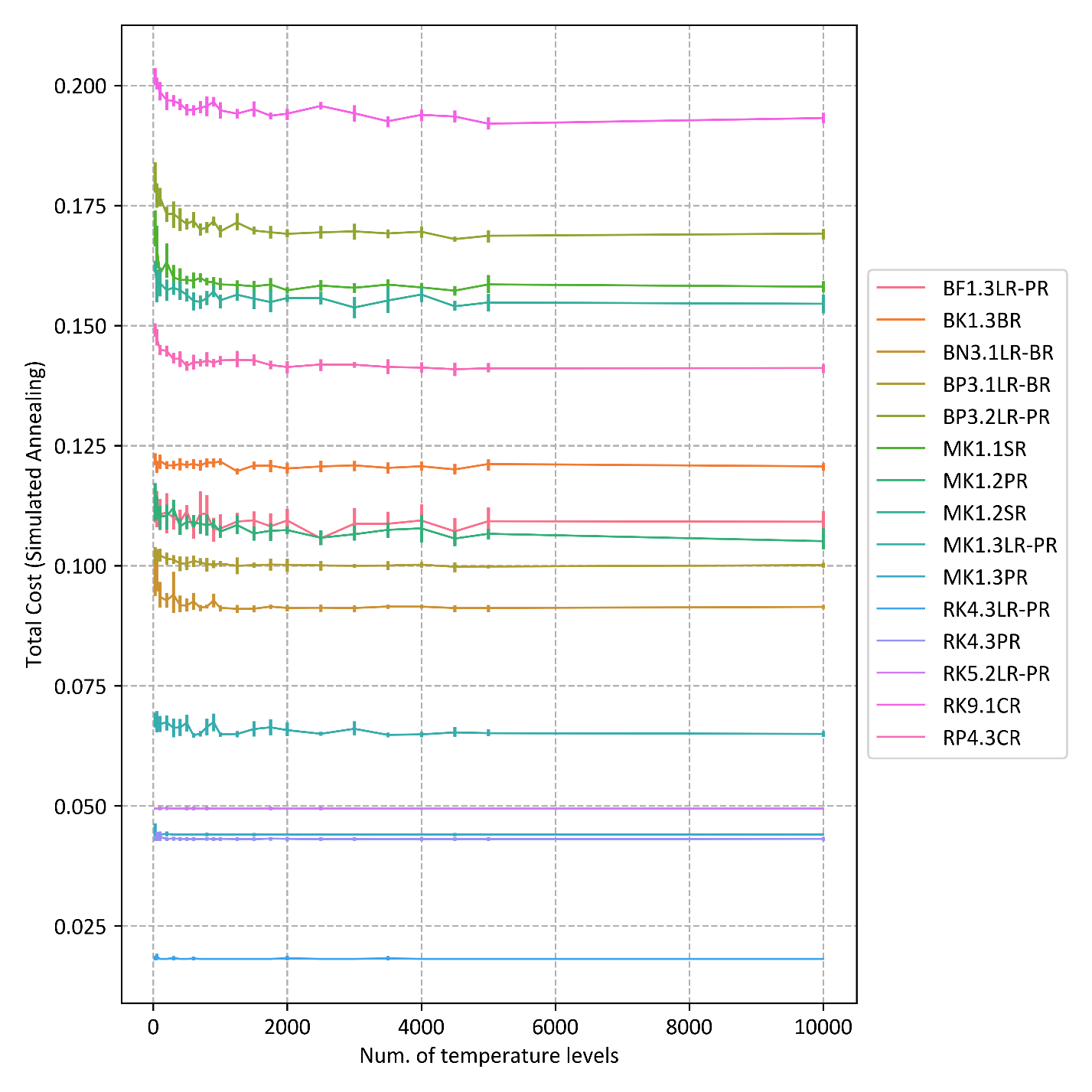

Figure S5: Total cost as a function of temperature levels for all fifteen images in Dataset Mahidol I. Each colored line represents results for one image. Measurements were taken for 25, 50, 100, 200, 300, 400, 500, 600, 700, 800, 900, 1000, 1250, 1500, 1750, 2000, 2500, 3000, 3500, 4000, 4500, 5000, and 10000 temperature levels. Results were computed five times for each image and at each level. The line indicates the corresponding mean values and the vertical bars the 95% confidence interval.

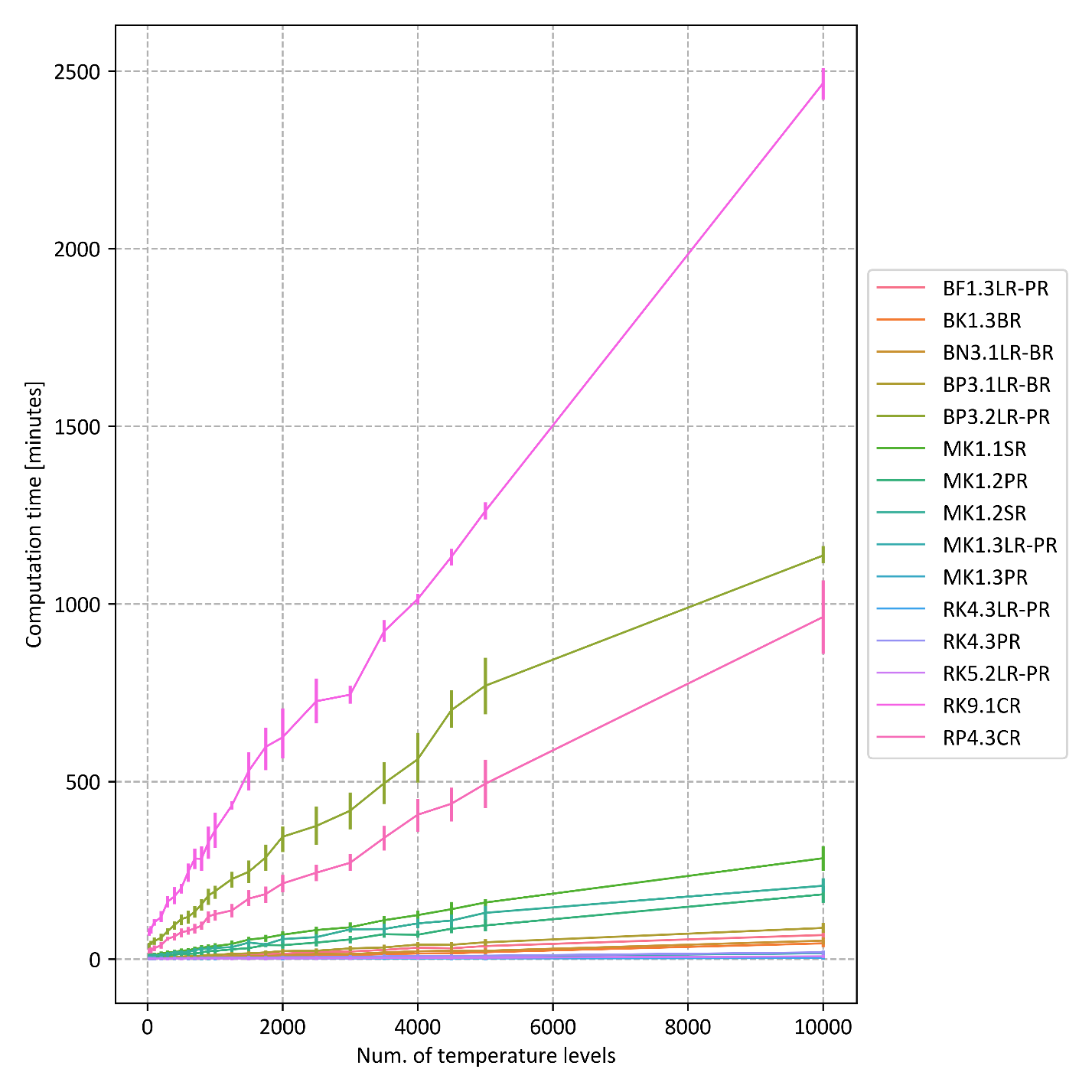

Figure S6: Computation time as a function of temperature levels for all fifteen images in Dataset Mahidol I. Each colored line represents results for one image. Measurements were taken for 25, 50, 100, 200, 300, 400, 500, 600, 700, 800, 900, 1000, 1250, 1500, 1750, 2000, 2500, 3000, 3500, 4000, 4500, 5000, and 10000 temperature levels. Results were computed five times for each image and at each level. The line indicates the corresponding mean values and the vertical bars the 95% confidence interval.

#### 1.2 Weight sensitivity

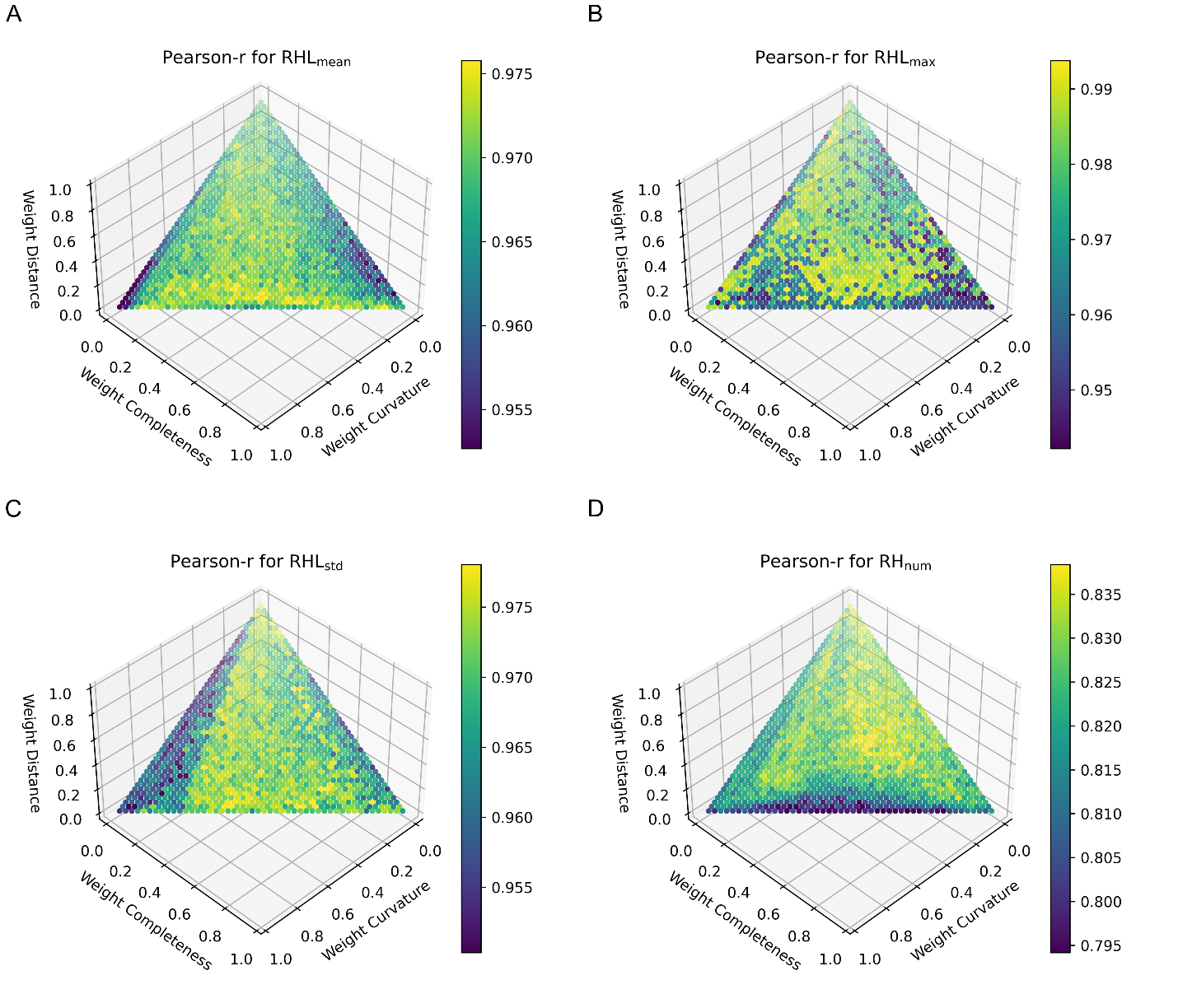

Figure S7: Pearson correlation coefficients indicated by color between manual and automatic measurements for different values of simulated annealing weights. Correlations are shown for four measured variables: (A) Correlation coefficient for RHL_mean_, (B) correlation coefficient for RHL_max_, (C) correlation coefficient for RHL_std_, and (D) correlation coefficient for RH_num_.

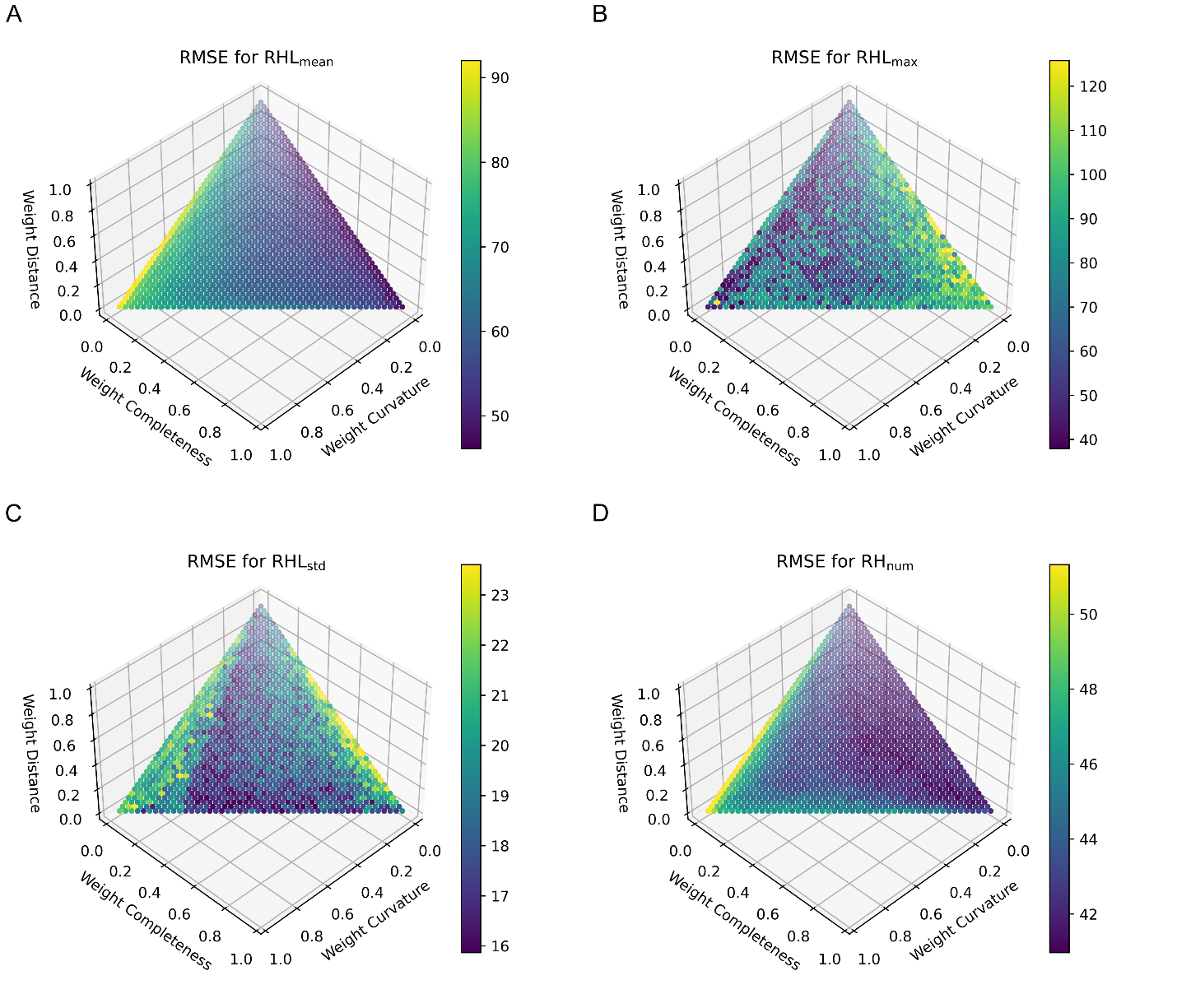

Figure S8: RMSE between manual and automatic measurements for different values of SA weights as indicated by color. Correlations are shown for four measured variables: (A) RMSE for RHL_mean_, (B) RMSE for RHL_max_, (C) RMSE for RHL_std_, and (D) RMSE for RH_num_.

### Computational Time

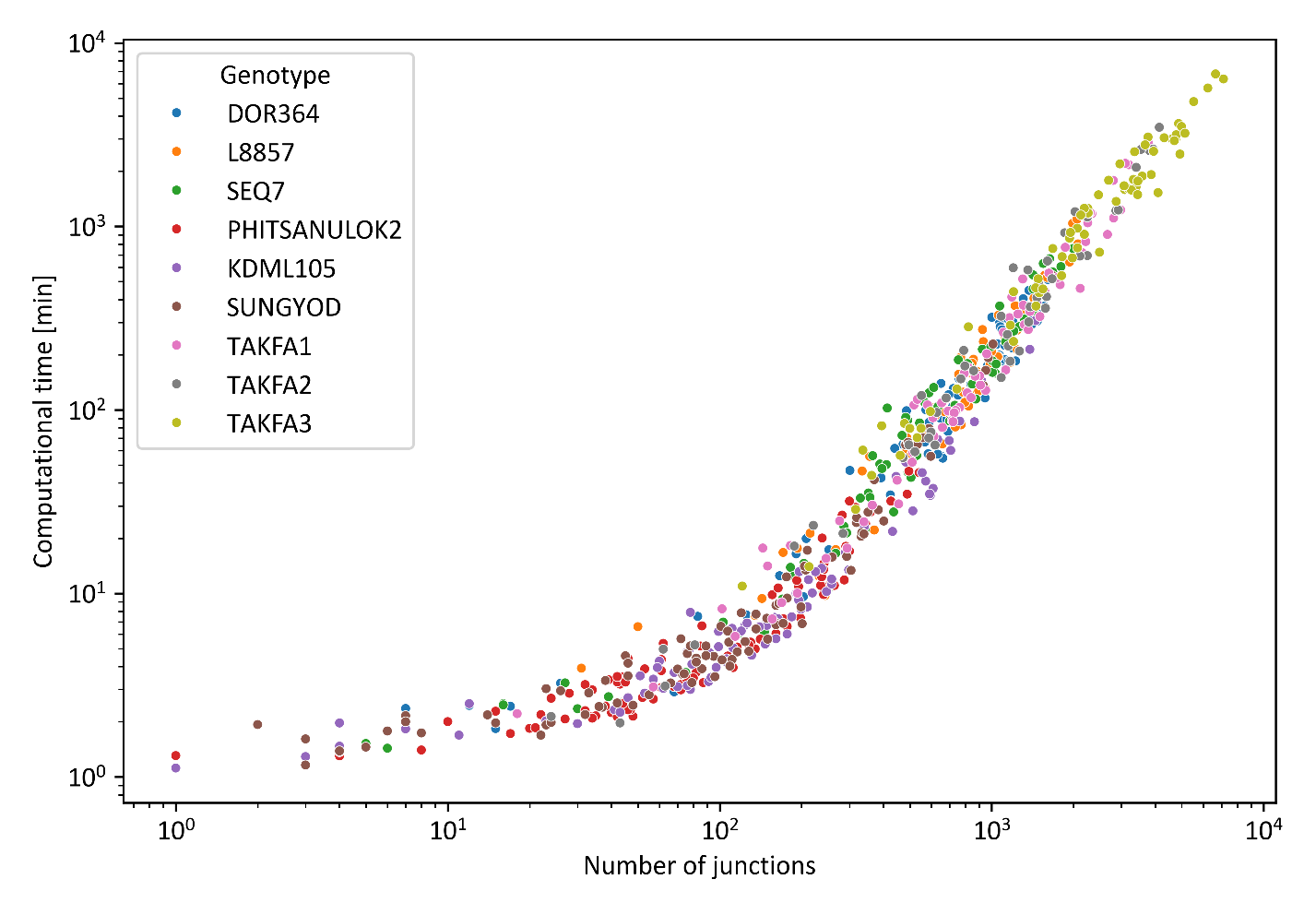

Figure S9: Computation time as function of junctions for images in Dataset Mahidol II. Each measurement point represents one image and shows the number of junctions on the medial axis and the time required to process the image in log10 scale. The color depicts the genotype.

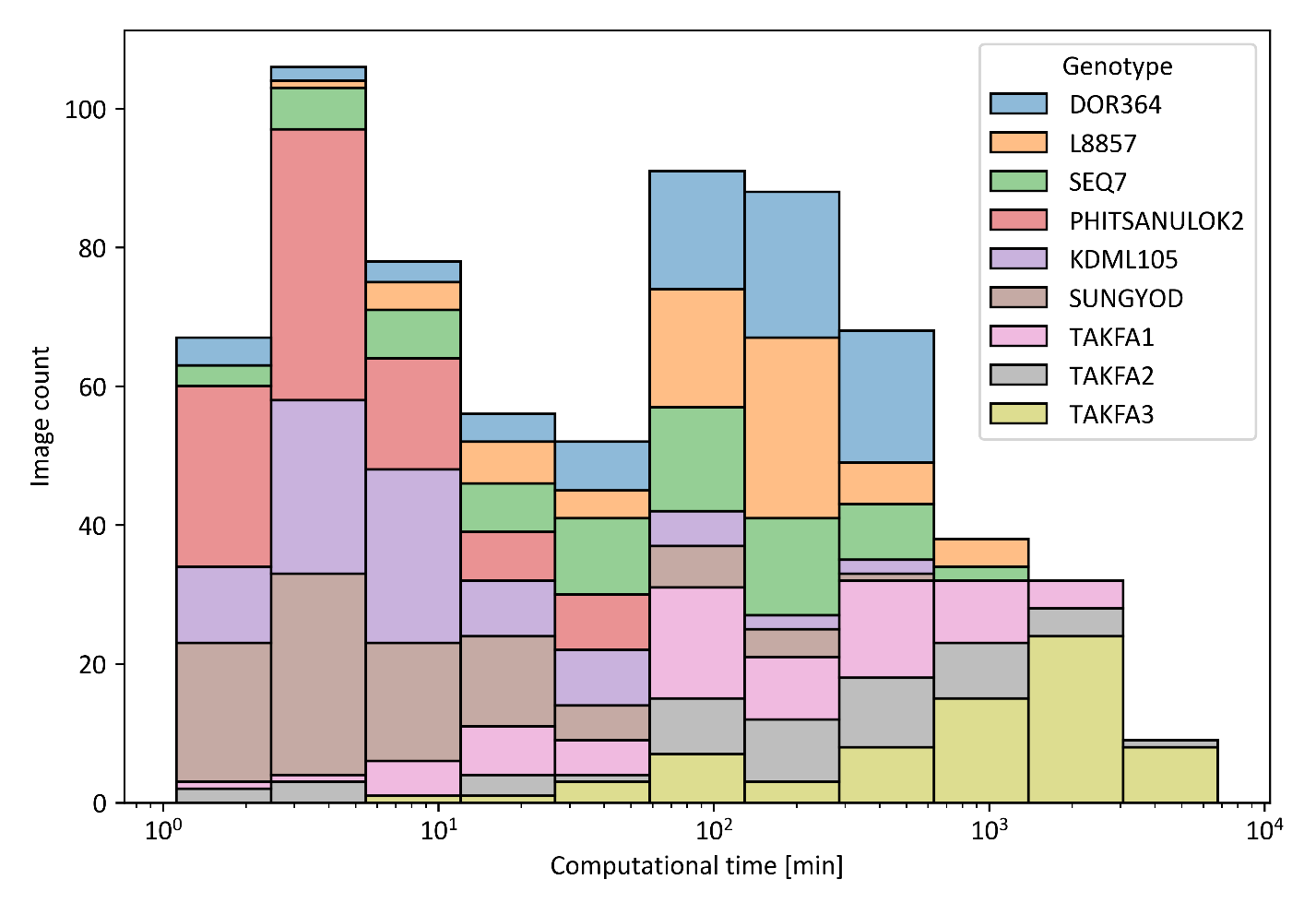

Figure S10: Histogram of computational time to process images in Dataset Mahidol I colored by genotype. The horizontal axis depicts the required computation time required to process an image and the vertical axis depicts the number of images falling into each time bin.

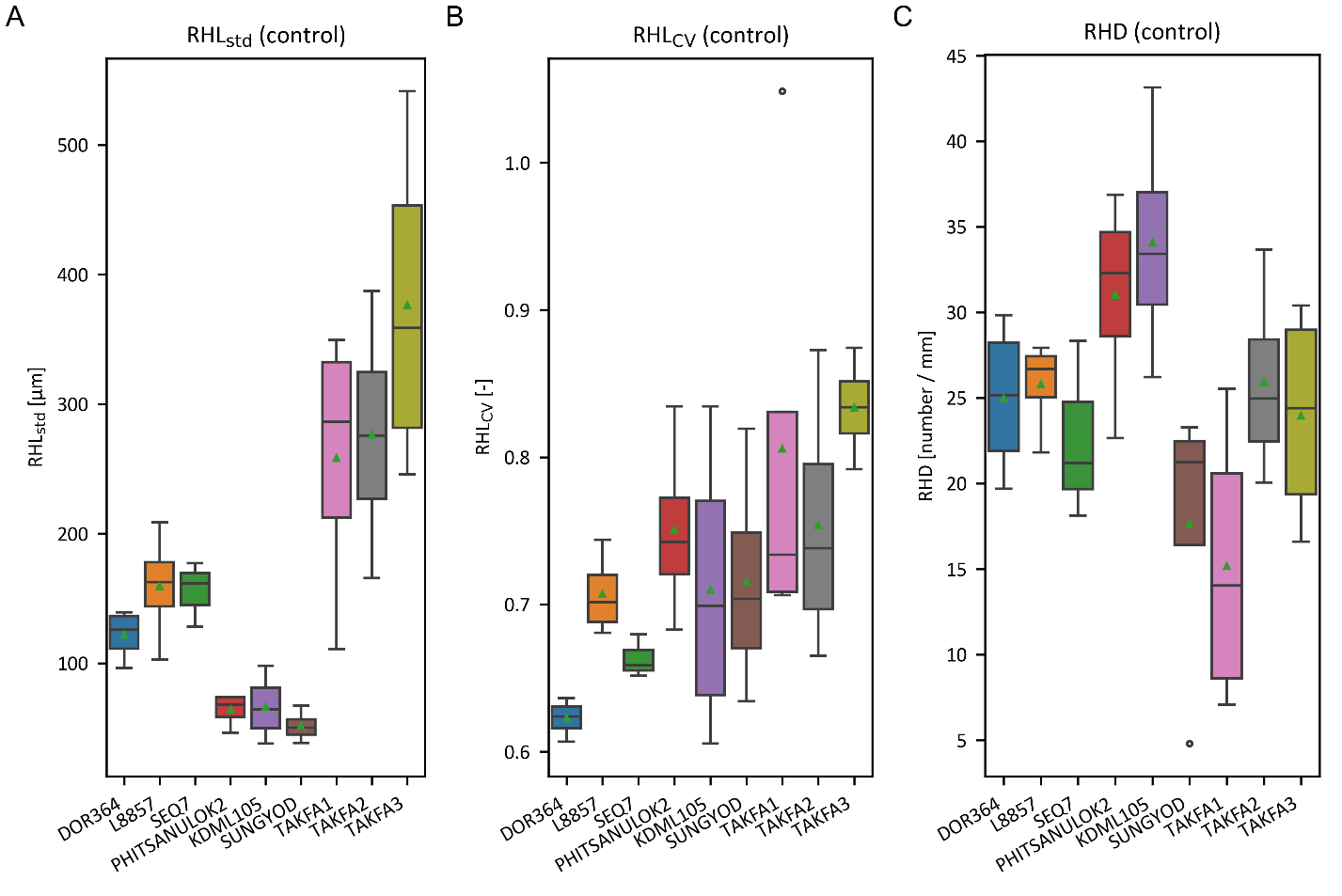

Figure S11: Measurements of root hair traits in the control groups of Dataset Mahidol II. The box displays the quartiles of the distribution, the middle horizontal line represents the median, and the whiskers extend to 1.5 times the IQR from the lower and upper quartiles. Outliers outside the whiskers are represented as circles. The mean of the distribution is displayed as a green triangle. (A) Distribution of RHL_std_ computed in individual plants. (B) Distribution of RHL_CV_ computed in individual plants. (C) Distribution of RHD computed for individual plants.

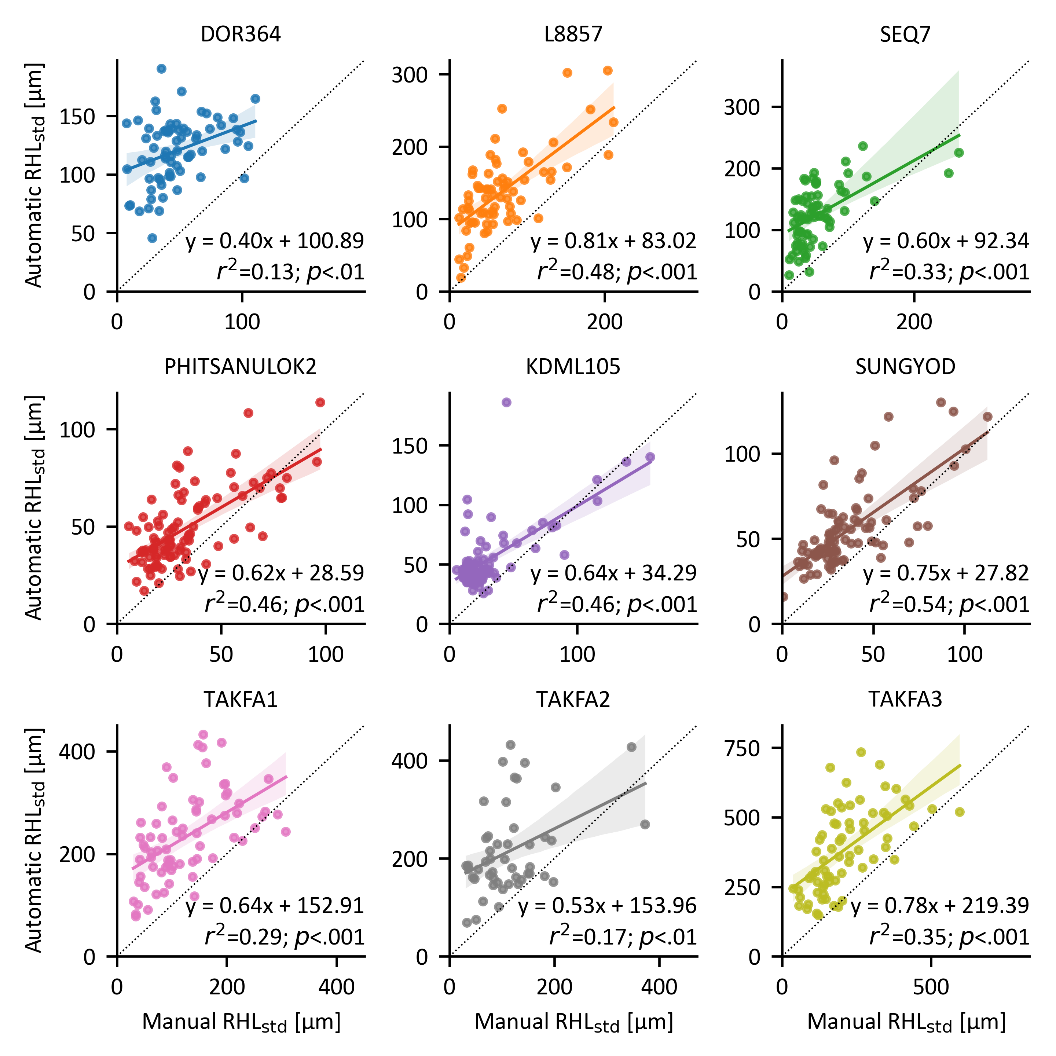

Figure S12: Correlation for automatic and manual measurements of RHL_std_. Each plot includes the results of a single genotype. Each circle represents RHL_std_ per image. The lines represent the linear regression models with 95% confidence intervals.

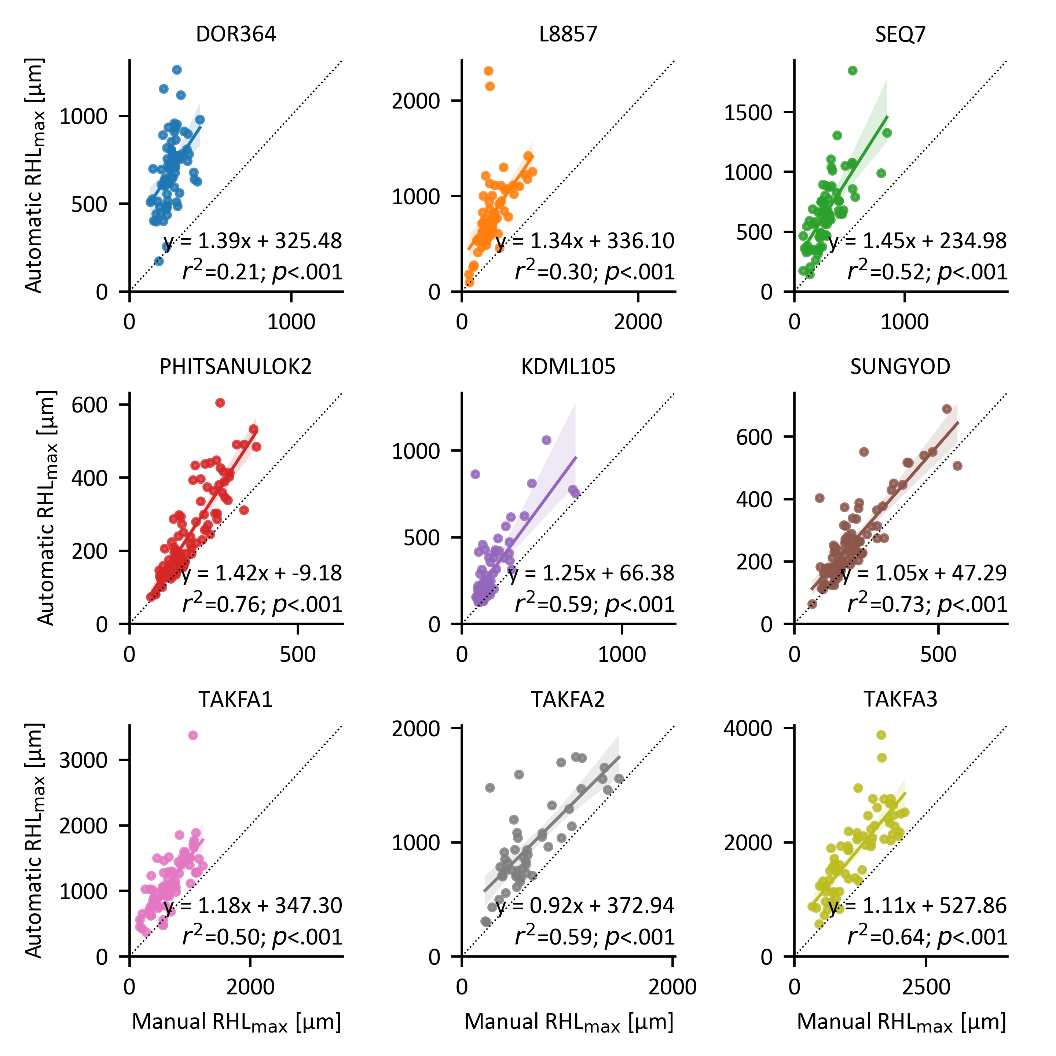

Figure S13: Correlation for automatic and manual measurements of RHL_max_. Each plot includes the results of a single genotype. Each circle represents RHL_max_ per image. The lines represent the linear regression models with 95% confidence intervals.

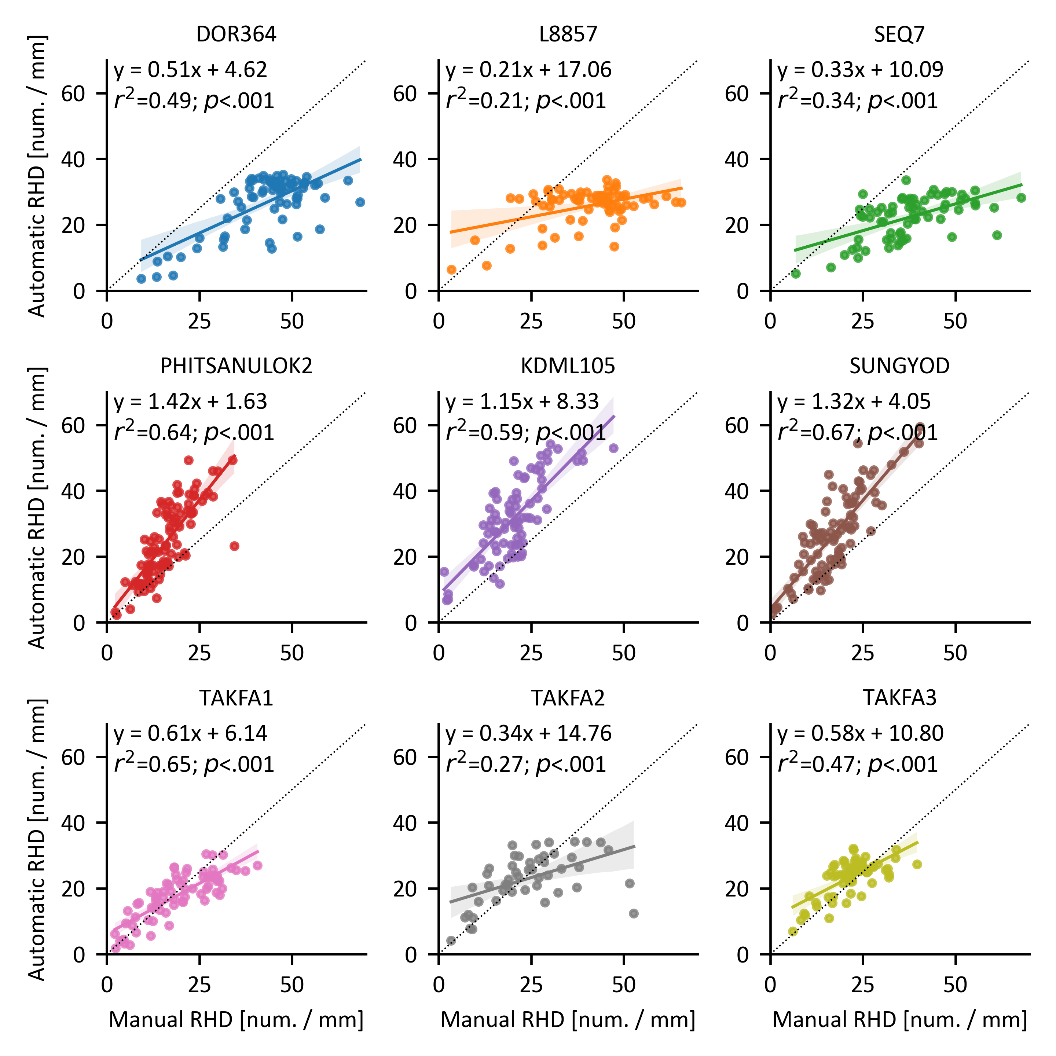

Figure S14: Correlation for automatic and manual measurements of RHD. Each plot includes the results of a single genotype. Each circle represents the RHD in a single image. The lines represent the linear regression models with 95% confidence intervals.
